# Stability of cross-sensory input to primary somatosensory cortex across experience

**DOI:** 10.1101/2024.08.07.607026

**Authors:** Daniel D Kato, Randy M Bruno

## Abstract

Merging information from across sensory modalities is key to forming robust, disambiguated percepts of the world, yet how the brain achieves this feat remains unclear. Recent observations of cross-modal influences in primary sensory cortical areas have suggested that multisensory integration may occur in the earliest stages of cortical processing, but the role of these responses is still poorly understood. We address these questions by testing several hypotheses about the possible functions served by auditory influences on the barrel field of mouse primary somatosensory cortex (S1) using *in vivo* 2-photon calcium imaging. We observed sound-evoked spiking activity in a small fraction of cells overall, and moreover that this sparse activity was insufficient to encode auditory stimulus identity; few cells responded preferentially to one sound or another, and a linear classifier trained to decode auditory stimuli from population activity performed barely above chance. Moreover S1 did not encode information about specific audio-tactile feature conjunctions that we tested. Our ability to decode auditory audio-tactile stimuli from neural activity remained unchanged after both passive experience and reinforcement. Collectively, these results suggest that while a primary sensory cortex is highly plastic with regard to its own modality, the influence of other modalities are remarkably stable and play a largely stimulus-non-specific role.

## Introduction

The mammalian cortex has traditionally been conceived of as comprising numerous discrete, cytologically- and functionally-defined areas serving distinct computational roles and coordinating with each other via long-range, hierarchical or recurrent interareal connections. Among these regions are the primary sensory cortical areas, which are the sites of the earliest stages of cortical information processing. According to the historical view, these areas operate as parallel modules dedicated to detecting low-level features from a single sensory modality each (Penfield & Boldrey 1937, Walzl & Woolsey 1946, Hubel & Wiesel 1959, Haberly & Price 1978). These areas were thought to then pass output to downstream midbrain structures like superior colliculus and association cortical areas like posterior parietal and prefrontal cortex, which would integrate information from across modalities into robust, disambiguated multisensory percepts of the world (Sprague & Meikle 1965, Bruce et al. 1981, Jay & Sparks 1984, Meredith & Stein 1984, Leichnetz 2001, Ernst & Bülthoff 2004, Barraclough et al. 2005, Schlack et al. 2005, Stein & Stanford 2008, Olcese et al. 2013, Raposo et al. 2012, Raposo et al. 2014, Nikbakht et al. 2018).

However, more recent literature has called for a reappraisal of such distinct parcellation (Wallace et al. 2004, Ghazanfar & Schroeder 2006, Liang et al. 2013). Numerous anatomical studies have shown that there exist direct, monosynaptic connections between primary sensory cortical areas (Budinger et al. 2009, Charbonneau et al. 2012, Iurilli et al. 2012, Stehberg et al. 2014, Henschke et al. 2015 , Godenzini et al. 2021), and complementary functional studies have demonstrated that presentation of a stimulus of nearly any sensory modality can modulate, and in some cases even drive, responses in nearly any primary sensory cortical area. For example, different types of auditory stimuli have been variously shown to hyperpolarize, suppress visual responses, sharpen visual tuning curves, or enhance spiking in primary visual cortex (Iurilli et al. 2012, Ibrahim et al. 2016, Meijer et al. 2017, Deneux et al. 2019, Knöpfel et al. 2019, Garner & Keller 2022) as well as suppress (Zhang et al. 2020) or enhance (Godenzini et al. 2021) responses to tactile stimuli in primary somatosensory cortex. Visual stimuli have been shown to reset the phase of local field potential oscillations in primary somatosensory cortex (Sieben et al. 2013) and drive spiking in infragranular layers of primary auditory cortex (Morrill & Hasenstaub 2018), tactile stimuli have been shown to hyperpolarize both primary visual and auditory cortices (Iurilli et al. 2012) and reset the phase of oscillations in auditory cortex (Lakatos et al. 2007), and olfactory stimuli have been found to bidirectionally modulate responses in primary somatosensory cortex (Renard et al. 2022).

Nevertheless, the computational roles subserved by these early cortical cross-sensory interactions remain enigmatic. Numerous potential functions have been suggested, including tuning curve sharpening (Ibrahim et al. 2016), pattern completion (Durup & Fessard 1935, Bakin & Weinberger 1990, Knöpfel et al. 2019), denoising (Ernst & Bülthoff 2004), cancellation of predictable inputs (Garner & Keller 2022), and nonlinear encoding of specific multisensory feature combinations (Deneux et al. 2019). Yet in many cases, very little is known about the most basic representational properties of these cross-sensory influences, such as whether they actually encode any sensory information about an area’s non-preferred modality *per se*; indeed, recent work suggests that many of these effects may be accounted for by signals related to stimulus-evoked movement, rather than by the stimuli themselves (Bimbard et al. 2023). Moreover, little is known about if and how these early cortical cross-sensory interactions are modified by experience; while both passive experience and reinforcement learning have repeatedly been shown to robustly affect responses to a primary sensory cortical area’s preferred modality (Shuler et al. 2006, Pantoja et al. 2007, Pleger et al. 2008, Weis et al. 2013, Kato et al. 2015, Poort et al. 2015, Keller et al. 2017, Henschke et al. 2020, Rabinovich et al. 2022, Benezra et al. bioRxiv), it remains unclear whether these effects extend to non-preferred modalities as well. Thus, many questions about putative early-cortical multisensory integration remain.

In this study, we addressed these issues by directly testing several hypotheses about the representational properties of auditory influences on the barrel field of primary somatosensory cortex (S1). We found that when cochlear and behavioral responses to auditory stimuli were intensity-matched, somatosensory cortex encoded little to no information about auditory stimulus identity. Moreover, we found no significant evidence of nonlinear encoding of specific audio-tactile feature conjunctions in S1. Extramodal influences, when detected, were remarkably stable over the course of both passive experience and reinforcement with reward. This stability suggests that these effects are undergirded by qualitatively different types of synaptic inputs from those responsible for a primary sensory cortical area’s responses to its preferred modality, which are in contrast known to be highly plastic over the course of learning and experience.

## Results

### Auditory stimuli evoke responses in an extremely small fraction of S1 cells

We first sought to address whether auditory stimuli evoke activity in L2/3 pyramidal cells. To test this, we performed *in vivo* 2-photon calcium imaging in the barrel cortex of awake, head-fixed mice while delivering randomly interleaved auditory and tactile stimuli. Tactile whisker stimuli (‘W’) consisted of a stimulus pole driven through the whisker field by a stepper motor, deflecting the whiskers at an angular velocity of ∼1800°/sec. Auditory stimuli consisted of 2 different 300 ms band-limited noise bursts: one ranging from 8.5-10.5 KHz (‘N1’) and the other ranging from 16.5-18.5 KHz (‘N2’; see Fig. 1a). The location of barrel cortex in each mouse was verified using intrinsic signal optical imaging, and a suitable 2-photon imaging site was found within barrel cortex by registering surface vasculature across 2-photon and widefield fluorescence images (Figs. 1b, c). The locations of imaging sites in barrel cortex were further confirmed by the presence of whisker-evoked activity transients in ΔF/F time series for individually-segmented cells (Fig. 1d).

**Figure 1:**
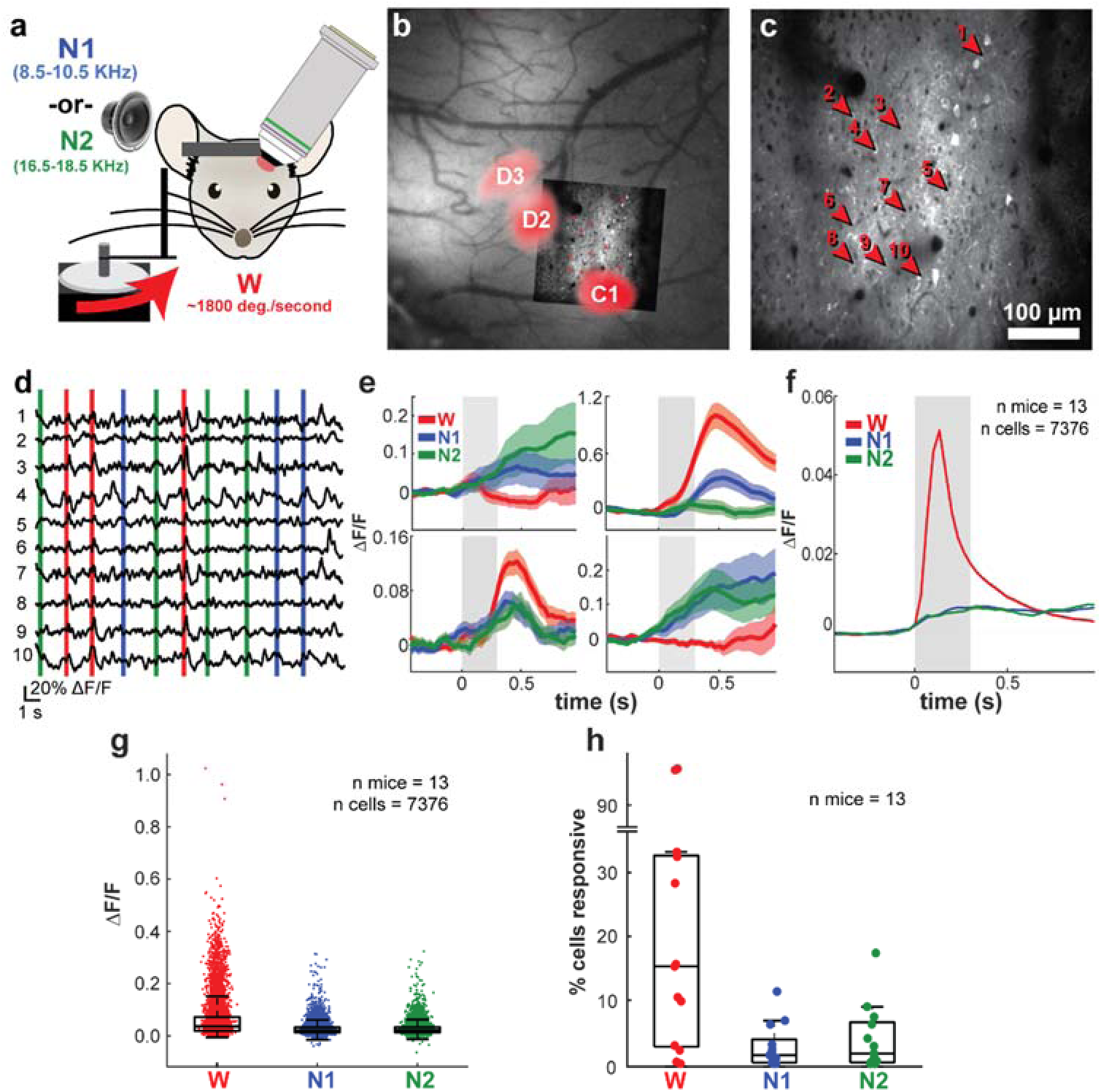
Auditory stimuli evoke responses in an extremely small fraction of S1 cells. **a**, Stimulus delivery and 2-photon imaging apparatus. **b**, Intrinsic signal map of barrel field of primary somatosensory cortex. Colored ellipses highlight locations of C1, D2, and D3 barrels. Inset: 2-photon field of view registered to surface vasculature in widefield image. **c**, Example 2-photon field-of-view, same as inset in **b**. Numbered arrowheads correspond to traces in **d**. **d**, Example ΔF/F time series from highlighted cells in **c**. Vertical bars represent stimulus onset times, color coded by condition as in **a**. **e**, Example trial-averaged responses of four neurons to a whisker stimulus alone, auditory stimulus 1 alone, or auditory stimulus 2 alone. Shaded area is standard error of the mean across trials. Gray rectangle: auditory stimulus epoch. **f**, Population-averaged responses to W, N1, and N2. Shaded area is standard error of the mean across 7,376 neurons. **g**, Trial-averaged ΔF/F response amplitude to W, N1, and N2 for all neurons across 13 mice. Box plots depict 25th, 50th, and 75th percentiles, whiskers depict 1.5 times interquartile range. **h** Percent cells significantly responsive to W, N1, and N2; each point one mouse and one condition.

Neurons exhibited heterogeneous responses to tactile and auditory stimuli. In addition to cells exhibiting classic whisker responses, some cells responded to both whisker and auditory stimuli, while others responded to auditory stimuli only (Fig. 1e). More generally, auditory stimuli were followed by small but sustained elevations in population-averaged activity (Fig. 1f). Compared to whisker responses, however, auditory response amplitudes were smaller and less variable. Across 7,376 neurons, trial-averaged responses to both N1 and N2 rarely exceeded 10% ΔF/F, with a median of about 2.1% ΔF/F and interquartile range of 1.8 percentage points (though note that these means include many response failures, and successful transients were often much larger); by contrast, the distribution of trial-averaged responses to whisker stimulation had a median of 3.5% ΔF/F and an interquartile range of about 4.9 percentage points, with about 8.3% of cells reaching response amplitudes between 15% and 30% ΔF/F (Fig. 1g). Statistically significant responses to both N1 and N2 were comparatively rare as well; across mice, the median percentage of cells significantly modulated by auditory stimulus onset (assessed by separately comparing each cell’s ΔF/F activity before and after stimulus onset, and corrected for multiple comparisons using false discovery rate procedures) was about 1.5% for both N1 and N2, although both stimuli evoked significant responses in more than 10% in some mice (Fig. 1h). By contrast, tactile stimuli evoked significant responses in approximately 15% of cells on average, consistent with previous results (O’Connor et al. 2010, Peron et al. 2015, Rodgers et al. 2021, Rabinovich et al. 2022).

### Barrel cortex weakly encodes auditory stimulus identity

Although we only observed weak responses to auditory stimuli in S1, the precise nature and quantity of the information encoded by these responses remained unclear. While these sound-evoked responses may encode genuine auditory sensory information, they may alternatively be driven by ancillary factors correlated with the presentation of an auditory stimulus, for example movement or arousal changes induced by the stimulus (Musall et al. 2019, Petty et al. 2021, Bimbard et al. 2023), or changes in some kind of internal arousal state (Allen et al. 2019).

We sought to address this question by testing whether barrel cortex differentially responds to or encodes information about the two auditory stimuli. If putative sound-evoked responses are actually driven by arousal state changes or sensory-evoked movements, and if N1 and N2 evoke sufficiently similar arousal state changes and/or movements, then they should encode little to no information about auditory stimulus identity *per se*. There also exists a spectrum of intermediate possibilities where some sound-evoked activity in S1 encodes genuine auditory sensory information, while other activity is driven instead by correlated variables like arousal state and movement.

In order to ensure that the overall intensity and salience of our auditory stimuli were well-matched, we performed *post hoc* auditory brainstem response recordings in every mouse, verifying that responses to N1 and N2 were nearly identical at the level of the cochlea and brainstem (Figs. 2a, b). Moreover, in order to confirm that N1 and N2 evoked similar behavioral responses, we recorded video of the whisker pad over the course of each imaging session and measured median whisker angle at each frame, providing a metric of overall whisker movement on each trial. We found that in a majority of mice (8/13), trial-averaged whisking responses to N1 and N2 were statistically indistinguishable (cf. example in top panel of Fig. 2c, Fig. S1a-i). In the remaining 5 mice, trial-averaged whisker angle during N1 and N2 presentation differed by at most only about 2-5 degrees (cf. example in Fig. 2c, bottom panel, Fig. S1j-m); in one of these mice, moreover, population-averaged S1 responses to N1 and N2 were nearly identical (Fig. S1n), meaning that differences in whisking behavior were not simply driving different overall levels of S1 activity in response to N1 and N2. Moreover, our previous results have shown that whisking is not a major driver of activity in barrel field L2/3 pyramidal cells (Rabinovich et al. 2022) and that firing rate changes of excitatory L2/3 neurons are only observed during large movements of nearly 50 degrees and then only by about 10% (Rodgers et al. 2021). These considerations taken together suggest that N1 and N2 are sufficiently behaviorally well-matched that any subtle difference in motor responses would be unlikely to explain auditory stimulus-specific activity in barrel cortex.

**Figure 2:**
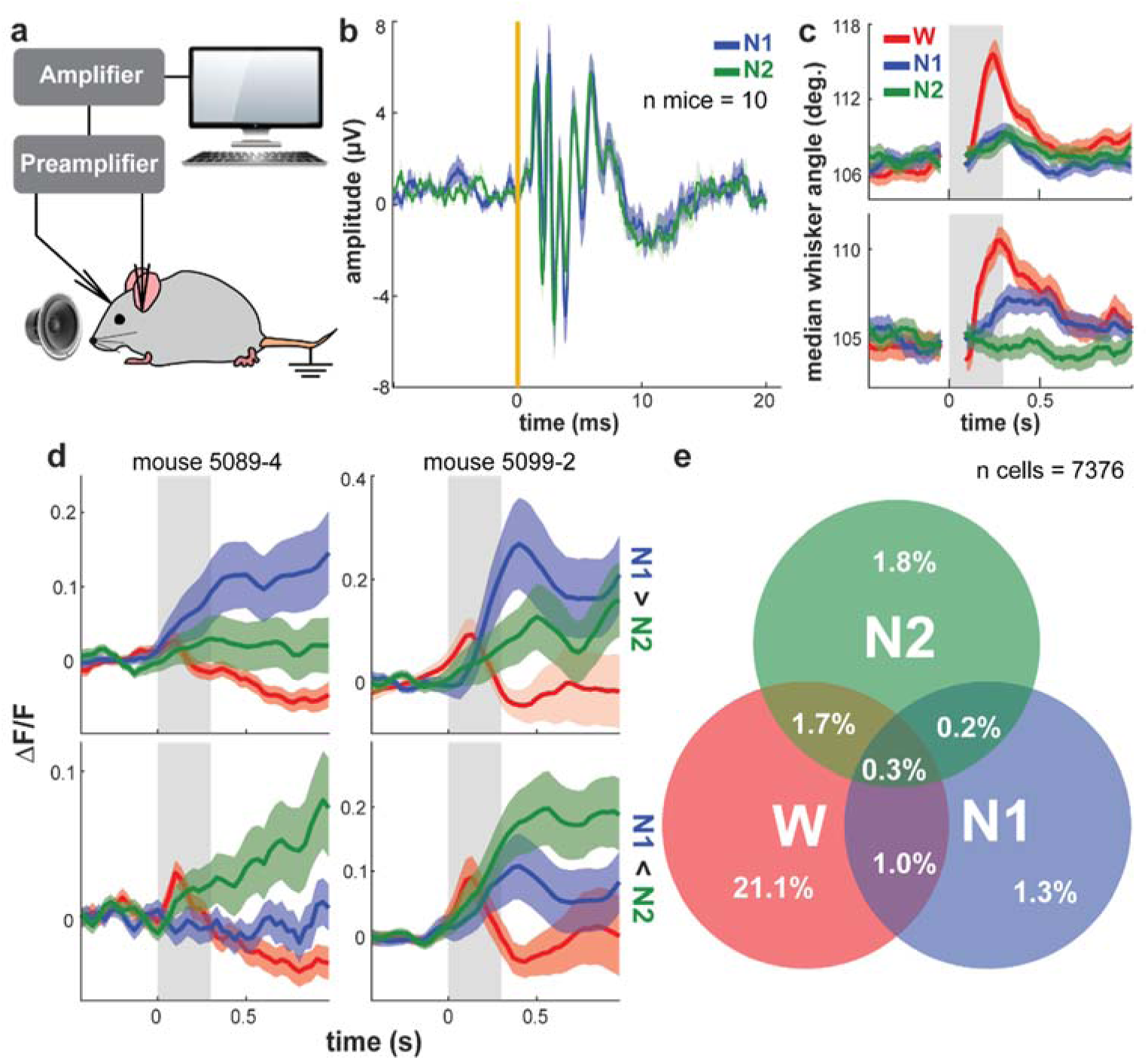
An extremely small fraction of S1 cells are auditory stimulus-selective. **a**, Auditory brainstem response recording setup. **b**, Mean auditory brainstem response to N1 and N2. Shaded area is standard error of the mean across mice. Yellow line: Stimulus onset time. **c**, Trial-averaged median whisker angle responses to W, N1, and N2 for two example mice. Shaded area is standard error of the mean across ∼100 trials per condition. **d**, Trial-averaged responses for four example neurons to W, N2, and N2. Top row: Neurons with higher mean response to N1 than to N2. Bottom row: neurons with higher mean response to N2 than to N1. Left column: Neurons from mouse 5089-4. Right column: Neurons from mouse 5099-2. **e**, Venn diagram of percentage of cells significantly responsive to W, N1, N2, and all combinations thereof, across 13 mice.

At the level of single cells, it was possible to find examples that preferentially responded to one auditory stimulus or the other. It was even possible to find cells with opposite auditory preferences in the same mouse and imaging site (Fig. 2d; note average whisking responses to N1 and N2 were nearly identical in these mice). While sound-responsive cells were rare overall, the majority of such cells were significantly responsive to only one auditory stimulus or the other; about 2.5% of cells preferred N1, 3.5% preferred N2, and only 0.5% of cells were significantly responsive to both (Fig. 2e; false discovery rate corrected). By contrast, nearly 20% of cells responded significantly only to the whisker stimulus.

Note, however, that the analyses discussed thus far consider each neuron singly, in contrast to any biologically plausible downstream readout cell or area, which would almost certainly pool input across many S1 neurons. Moreover, these single-cell analyses measure differences in trial-averaged responses over many stimulus presentations, which no downstream readout cell or area has access to at any given time. Thus, a more direct approach to quantifying the amount of auditory stimulus information available in S1 would be to consider how well auditory stimulus identity can be decoded on a trial-to-trial basis by a downstream readout pooling input across multiple barrel cortex cells. In order to account for these considerations, we trained a support vector machine (SVM) to decode auditory stimulus identity from neural population activity. Linear decoder performance was low, peaking marginally above chance (54.9%, Fig. 3a; p=0.04, bootstrap test over 1000 shuffles) at a single time bin around 800 ms after stimulus onset. Moreover, qualitatively similar results were obtained using a multilayer perceptron (MLP) nonlinear decoder (Fig. S2). These results collectively imply that S1 encodes a small amount of information about auditory stimulus identity, although trial-to-trial variability makes decoding highly unreliable. Consistent with this interpretation, the majority (∼70%) of sound-responsive cells exhibited significant ΔF/F transients only on a minority (∼10%) of trials (Fig. 3b).

**Figure 3:**
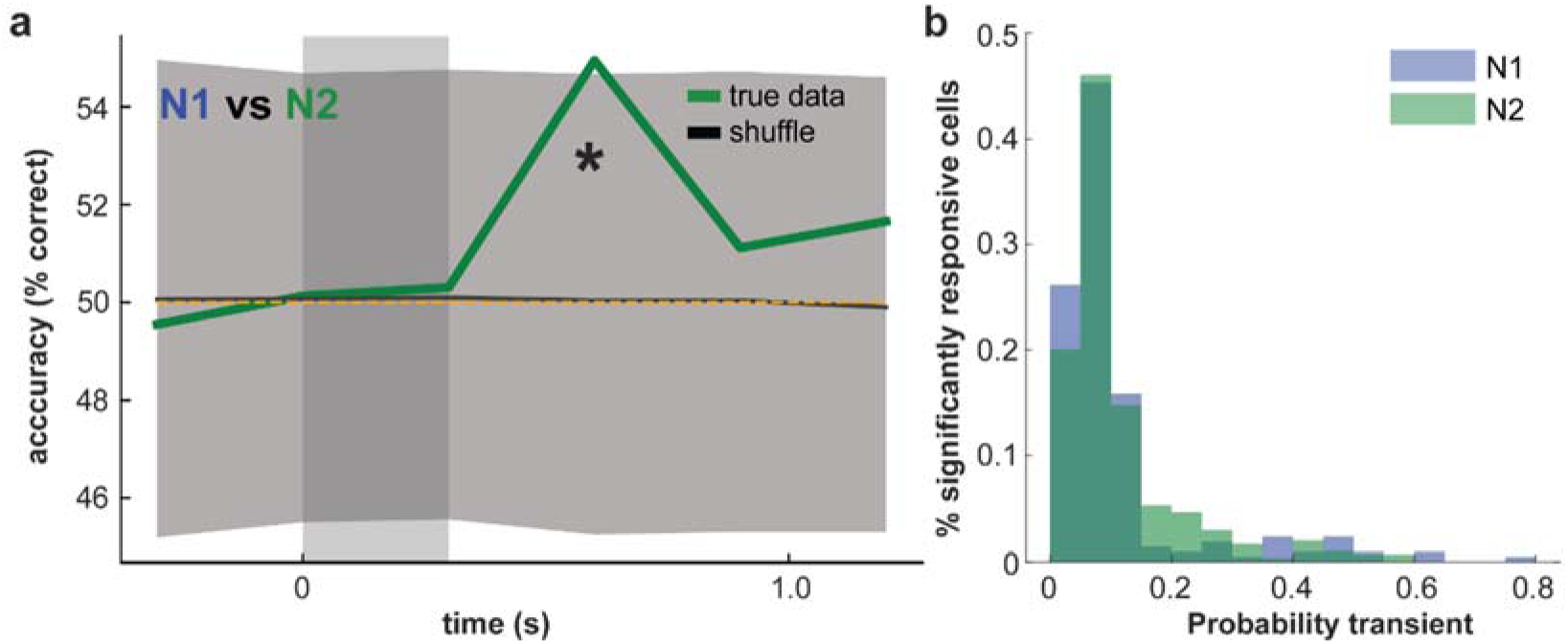
S1 weakly encodes auditory stimulus identity. **a**, Cross-validated N1 vs N2 SVM accuracy, bin size equals 400 ms. Green trace is mean cross-validated SVM accuracy over 50 random instances of a pseudosimultaneous imaging session across 13 mice and 7,376 neurons plus associated neuropil regions-of-interest for each. Gray shaded area is 95% confidence interval of shuffle distribution, n shuffles=1000. Vertical gray rectangle: auditory stimulus epoch. Yellow dotted line: chance accuracy level. Asterisk denotes time bin where accuracy is significantly different from chance. **b**, Histogram of probability of transient on N1 trials for N1-responsive cells (blue, 214 cells) and probability of transient on N2 trials for N2-responsive cells N2 (green, 304 cells).

### Auditory input weakly suppresses tactile responses in S1

Having thus found that S1 only weakly encodes information about pure auditory stimuli, we then went on to ask whether S1 might encode information about conjunctions of simultaneous or near-simultaneous auditory and tactile stimuli. It could be that previously described multisensory inputs to primary sensory cortical areas serve mainly to modulate responses to the dominant sensory modality, rather than to drive spiking responses on their own (Ibrahim et al. 2016, Zhang et al. 2020). For example, inputs conveying auditory signals could terminate onto distal apical dendrites of L2/3 S1 pyramidal cells, where, despite being too weak to strongly drive spiking activity on their own, they may interact with whisker-related inputs arriving through basal dendrites, enhancing or suppressing spiking responses to tactile stimuli (Fig. 4a), analogous to mechanisms that have previously been described in barrel cortex (Xu et al. 2012).

**Figure 4:**
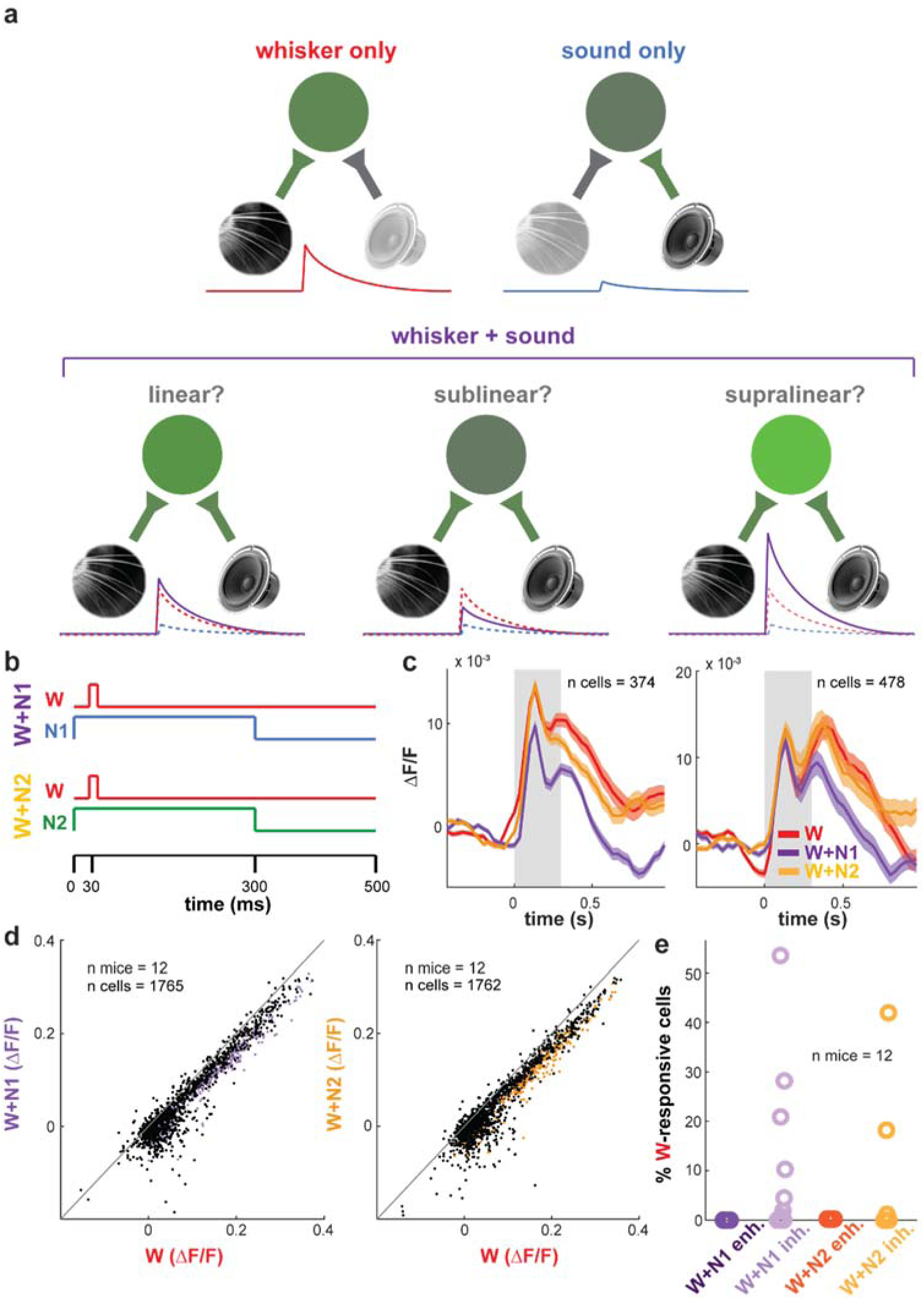
Auditory inputs weakly suppress tactile responses in S1. **a**, Schematic illustration of hypothetical forms of nonlinear audio-tactile integration in S1. Top left: whisker input by itself moderately drives activity in S1 neurons. Top right: auditory input by itself weakly drives activity in S1 neurons. Bottom left: linear summation, in which the S1 response to simultaneous whisker and sound (purple line) equals the arithmetic sum of responses to whisker and auditory stimuli alone (dashed red and blue lines, respectively). Bottom center: in sublinear summation, auditory input suppresses tactile responses, yielding lower responses than to the sum of whisker and auditory stimuli alone. Bottom right: in supralinear summation, auditory input enhances tactile responses, yielding higher responses than to the sum of whisker and auditory stimuli alone. **b**, Schematic illustration of individual trial structure. **c**, Population-averaged responses to W, W+N1(purple), and W+N2 (amber) for two example mice. Shaded area is standard error of the mean across neurons. **d**, Scatterplots of trial-average response AUC for conjunctive versus whisker-alone trials. Left panel: W+N1 vs W. Right panel: W+N2 vs W. Colored points denote cells with significantly different responses for W and conjunctive stimulus for each plot. **e**, Percent W-responsive cells significantly enhanced or inhibited by simultaneous presentation of N1 and N2.

In order to address this question, we introduced two additional stimulus conditions: whisker stimuli presented in conjunction with N1 (‘W+N1’), and whisker stimuli presented in conjunction with N2 (‘W+N2’). In both conditions, the speaker and motor turned on simultaneously, resulting in the stimulus pole hitting the whiskers on average about 30 ms after sound onset (Fig. 4b; although note with some trial-to-trial variability due to movement of the whiskers). This average latency was chosen, based on previously described sound-evoked postsynaptic potentials, so that auditory and tactile inputs arrived in barrel cortex nearly simultaneously (Iurilli et al. 2012).

Consistent with previous findings, we found that presenting sound and whisker stimuli simultaneously had a predominantly inhibitory effect on responses to whisker touch (Zhang et al. 2020, Iurilli et al. 2012, Ibrahim et al. 2016). Furthermore, W responses in some mice were differentially inhibited by N1 and N2, resulting in small-but-significant differences in population-averaged responses to W+N1 and W+N2 (Fig. 4c). In general, however, N1 and N2 elicited similar amounts of suppression (Fig. 4d). Among cells with significant responses to W, peak trial-averaged responses were significantly lower for both W+N1 and W+N2 (signed rank tests, Z=34.38, p<0.0001 and Z=34.79, p<0.0001, respectively). N1 and N2 both significantly inhibited total of about 13% of W-responsive cells, although this figure varied considerably between mice; while <5% of W-responsive cells were significantly inhibited by sound in most mice, this figure reached up to 40-50% in other mice (Fig. 4e). Notably, no cells showed significant enhancement of whisker responses by either sound. Consistent with the finding that sound has a weakly suppressive effect on whisker touch responses in barrel cortex, we found that a linear decoder was able to classify whisker-alone versus whisker-plus-sound trials with low but significantly above-chance accuracy (Fig. 5b; p=0.009, bootstrap test over 1000 shuffles).

**Figure 5:**
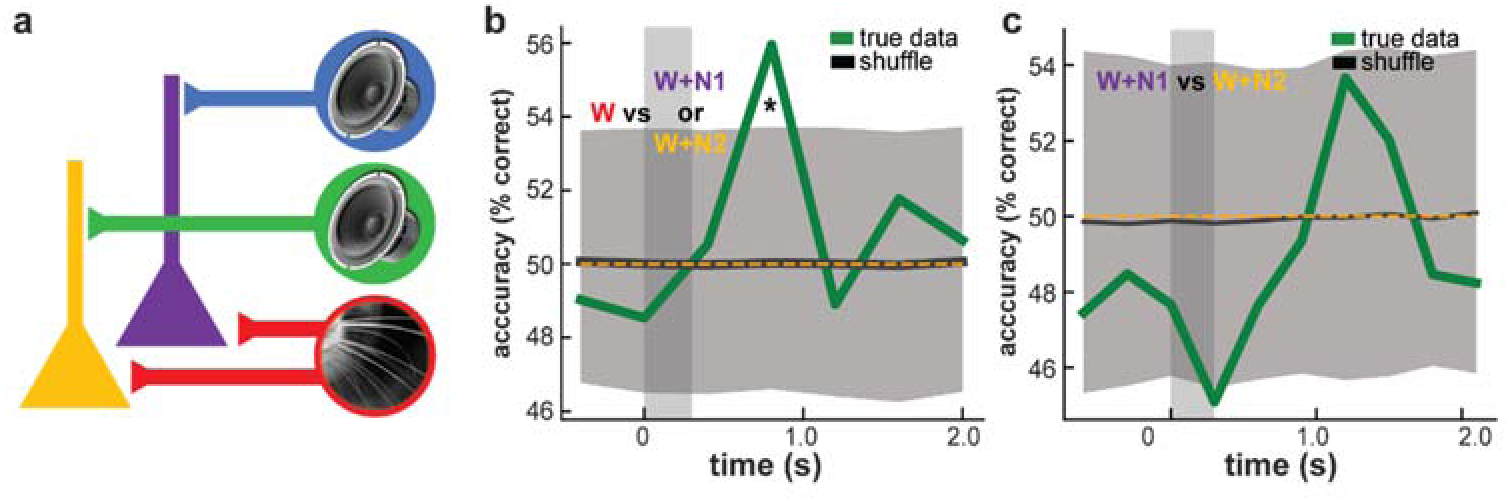
S1 does not encode audio-tactile stimulus identity. **a**, Schematic illustration of nonlinear mixed selectivity. In the purple cell, responses to input from W (red) are selectively modulated by input from N1 (blue); in the yellow cell, W responses are selectively modulated by input from N2(green). **b**, SVM accuracy for W vs conjunctive stimulus trials (i.e., one class including trials of type W, the other including trials of both types W+N1and W+N2), bin size equals 400 ms. Green trace: mean SVM accuracy over 50 pseudosession instances generated from 13 mice and 7,376 neurons plus associated neuropil regions-of-interest for each, gray shaded area: 95% confidence interval of accuracy distribution over 1000 shuffles. **c**, Same as **b**, but for classifying W+N1vs W+N2. Mice and neurons are the same as in Fig. 3.

We then considered the possibility that this suppressive modulatory input could be distributed across the S1 population in an auditory stimulus-specific manner, with different whisker-responsive cells selectively suppressed by different auditory stimuli. Such sound-specific modulation would effectively encode audio-tactile stimulus identity, as different sounds presented concurrently with the same tactile stimulus would result in different patterns of suppression of the tactile response (Fig. 5a). Note that any such encoding of audio-tactile stimulus identity would indicate nonlinear mixing at the spiking level, since linearly summing tactile responses with the largely auditory stimulus non-specific spiking responses already observed in S1 would not encode any further information about auditory stimulus identity, whether singly or in conjunction with a concurrent tactile stimulus. Indeed, such nonlinear mixing has been repeatedly shown to be indispensable for flexibly performing complex behavioral tasks (Rigotti et al. 2013, Fusi et al. 2016, Bernardi et al. 2020, Nogueira et al. 2023). We sought to quantify how much information about audio-tactile stimulus identity is available in S1 on a trial-by-trial basis by training an SVM to classify W+N1 vs W+N2 trials, finding that performance fell within chance levels (Fig. 5c; peak SVM performance = 53.6%, p=0.08).As with the pure auditory stimuli, training a nonlinear MLP to classify W+N1 vs W+N2 yielded qualitatively similar results to a linear decoder (Fig. S3). These results therefore support the conclusion that in naive mice, these sounds generally has a mild suppressive effect on whisker responses in S1, but does not reliably encode information about specific audio-tactile conjunctions on a trial-by-trial basis.

### Auditory information in S1 is stable over the course of passive experience

Our results thus far demonstrate that, in naive mice, sound-evoked responses in S1 encode little information about auditory stimulus identity and almost no information about audio-tactile stimulus identity conjunctions. Nevertheless, inputs conveying sound-evoked signals to S1 exist, though weak. We therefore sought to characterize whether these inputs are plastic, and whether various forms of experience might affect the information content of these inputs. We first considered a model in which Hebbian-like plasticity causes weak or latent auditory inputs to reactivate ensembles of S1 neurons encoding correlated whisker stimuli. In this model, if an auditory and tactile stimulus are sufficiently correlated, then as the firing of latent, sound-encoding presynaptic inputs becomes correlated with the firing of touch-encoding postsynaptic cells in S1, previously silent synapses undergo potentiation such that the auditory stimulus acquires the ability to reactivate the ensemble of S1 neurons encoding the correlated tactile input. Previous theoretical work has suggested that this kind of mutual reactivation of neuronal ensembles encoding correlated stimuli in different sensory areas could subserve cross-sensory coordinate transformations (Pouget et al. 2002).

To test this hypothesis, we subjected mice (n=4) to a passive pairing paradigm in which one auditory stimulus was repeatedly paired with a tactile stimulus over the course of several days (Fig. 6a). On the first day of the paradigm (‘pre-pairing phase’), mice were imaged while presented with randomly interleaved trials of 6 stimulus conditions: W, N1, N2, W+N1, W+N2, and a catch condition in which neither whisker nor auditory stimulus was presented. Over the course of the subsequent 3 days (‘pairing phase’), mice were imaged while repeatedly presented with only 2 of the 6 stimulus conditions: W+N1, and N2 alone. W and N1 were thus perfectly correlated during the pairing phase, and the inclusion of N2 alone trials ensured that total exposure to N1 and N2 (in isolation or in conjunction with W) was balanced across pairing to prevent any overall effects of statistical frequency like habituation, etc. Again, sound onset preceded whisker contact by on average about 30 ms (with some trial-to-trial variability due to whisker movement), designed to result in sound-evoked inputs arriving in S1 simultaneously with or a few milliseconds before tactile response onset. After pairing, S1 responses to all 6 stimulus conditions were probed during imaging once again (‘post-pairing’).

**Figure 6:**
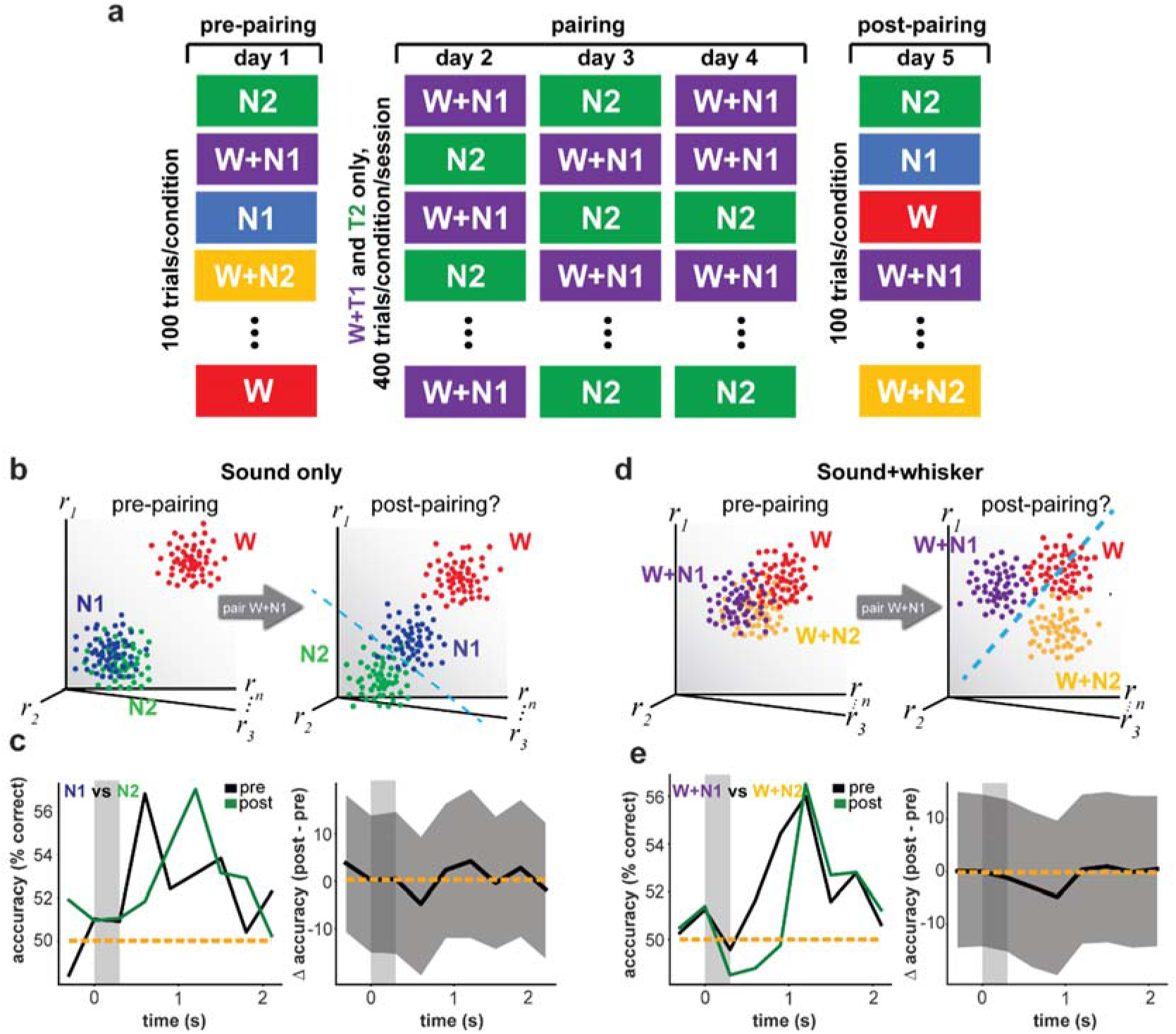
Auditory information in S1 is stable over passive experience. **a**, Diagram of pairing paradigm. **b**, Illustration of Hebbian audio-tactile association learning model prediction. Left panel: Before pairing, N1 and N2 evoke small responses across neurons (blue and green point clouds close to origin), while W evokes comparatively large responses across many neurons (red point cloud distant from origin). Right panel: Post-pairing, N1 reactivates many of the neurons activated by W, causing blue point cloud to shift towards red. Since N2 is unpaired with W and does not reactivate W-responsive cells, green point cloud remains near origin. Consequently, N1 and N2 point clouds diverge, become more decodable. **c**, Left panel: Mean pre-and post-pairing N1 vs N2 SVM accuracy for 1000 bootstrapped pairs of pre-and post-pairing pseudosessions over 4 mice (1,665 cells pre-pairing, 1,184 cells post-pairing), bin size equals 300 ms. Right panel: Post-minus pre-pairing N1 vs N2 SVM accuracy. Black trace: Mean difference over 1000 bootstrapped pairs of pre- and post-pairing pseudosession instances. Grey shaded area: 95% confidence interval of distribution of post-pre accuracy differences. **d**, Illustration of experience-dependent emergence of nonlinear mixed selectivity. Top panel: Before pairing, both cells are purely selective for the whisker stimulus. Bottom panel: Post-pairing, different cells develop nonlinear mixed selectivity for specific audio-tactile stimuli. **c**, Illustration of experience-dependent change in population audio-tactile responses. Top panel: Before pairing, W+N1and W+N2 are largely non-separable. Bottom panel: after pairing, W responses of different S1 subpopulations are differentially enhanced or suppressed by N1 and N2, leading to distinct, separable population responses to W+N1and W+N2. **e**, Same as **c**, but for W+N1vs W+N2.

In order to test for auditory stimulus-specific reactivation of correlated S1 subpopulations following experience, we compared N1 vs N2 SVM performance before and after pairing. If after pairing N1 comes to even partially reactivate the neuronal ensemble encoding W, then the population response to N1 should start to more closely resemble the population response to W. In other words, the population response to N1 should shift closer to the population response to W in neural state space (Fig. 6b). By contrast, because N2 was not correlated with W, then if auditory input to S1 supports the learning of specific audio-tactile correlations, N2 should not reactivate the ensemble encoding W, and the N2 population response should not shift closer to the W population response. Given that the N1 and N2 population responses are highly separable from the W population response to begin with, a selective shift of N1 towards W responses would cause a divergence between N1 and N2 population responses, leading to improved decodability for N1 and N2.

Before pairing, N1 vs N2 decoder performance was once again modest, peaking on average at 56.7% (Fig. 6c, left panel). Similarly, post-pairing SVM performance peaked at 56.9%, albeit in the subsequent time bin 400 ms later. A bootstrap test of differences between pre- and post-pairing SVM accuracy showed that the change in performance was not significantly different from 0 for any time bin, suggesting that overall amounts of information about auditory stimulus identity in S1 remained stable following passive experience (Fig. 6c, right panel; p>0.34, bootstrap test over 1000 iterations). Nonlinear decoders yielded qualitatively similar results, performing largely at chance levels throughout the trial epoch after pairing just as before pairing (Fig. S4). We thus find no evidence that passive experience enables auditory stimulus-specific reactivation of correlated touch-coding S1 ensembles via Hebbian-like plasticity under the present experimental conditions. Importantly, while our analyses do not rule out that associations may be learned elsewhere in the brain, or even at the behavioral level, our decoder results reveal no evidence of such learning occurring in S1 itself.

Having thus shown that information levels about pure auditory stimulus identity remain stable over the course of passive experience, we went on to ask whether conjunctive audio-tactile stimulus encoding is affected by passive experience. Previous experimental work has shown that appropriately timed input to proximal and distal dendrites of pyramidal cells can potentiate distal inputs, which interact nonlinearly with proximal inputs to drive burst firing in hippocampus (Takahashi et al. 2009). We thus considered whether S1 might be subject to a similar form of plasticity, whereby latent nonlinear interactions between auditory and tactile inputs must be potentiated through pairing before being expressed as strong modulation of whisker responses by a simultaneous auditory stimulus. If such plasticity took place in a stimulus-specific manner, it would provide a means for S1 cells to acquire selectivity to common audio-tactile feature conjunctions in the environment, resulting in highly distinct, separable representations in neural state space (Fig. 6d). We tested this possibility by measuring whether W+N1vs W+N2 decoder performance improved over the course of passive experience. We found again that decoder accuracy was modest, peaking on average around 56% both before and after pairing (Fig. 6e, left panel), and a bootstrap test confirmed that change in decoder performance was not significantly different from 0 (Fig. 6e, right panel, p>0.453). Similarly, nonlinear MLP performance showed little to no improvement after passive pairing, confined largely to chance levels throughout the trial epoch as before pairing (Fig. S5). These results thus demonstrate that as with information about pure auditory stimulus identity, the overall amount of information about audio-tactile stimulus conjunctions in S1 remains unchanged over the course of passive experience. Again, while our findings do not preclude the possibility that passive pairing alters the representation of audio-tactile stimuli elsewhere in the brain, our results reveal no evidence of such learning in S1 itself.

### Auditory information in S1 is stable over the course of reward conditioning

Having thus found that passive experience alone is insufficient to induce changes in the overall strength of pure auditory or audio-tactile stimulus encoding in S1, we considered whether pairing audio-tactile stimuli with reward drove plasticity. Previous results from our lab have shown that reward is necessary to enhance responses to tactile stimuli in S1 (Rabinovich et al. 2022, Benezra et al. under review), and analogous work in mouse primary visual cortex has shown that pairing 2 distinct visual stimuli with reward increases decoder performance (Henschke et al. 2020). Such findings suggest that auditory inputs to S1 could be governed by a “three-factor plasticity” rule, whereby latent auditory inputs onto postsynaptic cells with correlated whisker-evoked firing are only potentiated in the presence of some gating cue that signals behavioral relevance, like reward.

To test this idea, another cohort of mice (n=9) was subjected to a pairing paradigm in which audio-tactile stimuli predicted reward. As in the previous paradigm, the pre-pairing phase consisted of a single probe session in which mice received all 6 stimulus conditions. During pairing, mice received only W+N1 and W+N2 trials (Figs. 7a, b). In order to prevent different reward contingencies from driving differential changes in consummatory behavior-related activity that could mask changes in sensory representations *per se*, both stimuli were followed by a water reward after a 200 ms trace interval; rewarding both stimuli ensured that any behavioral or arousal state changes related to reward anticipation or consumption would be matched across the two trial conditions, and thus any improvement in decoder performance would have to be due to changes in sensory representations themselves. Crucially, moreover, note that previous research shows that pairing two different stimuli with reward equally does *not* cause their representations to overlap trivially due to similar behavioral responses; to the contrary, Henschke et al. (2020) show that pairing two visual stimuli with reward equally actually causes them to become *more* decodable in V1. Anticipatory licking during the trace interval was used as a metric of engagement and learning that audio-tactile stimuli cued reward. Mice were given 2 weeks of pairing, after which they were given an additional week of pairing if they had not anticipatorily licked on at least 70% of trials over the previous 2 days. After pairing, mice received an additional probe session consisting of all 6 conditions with no reward.

**Figure 7:**
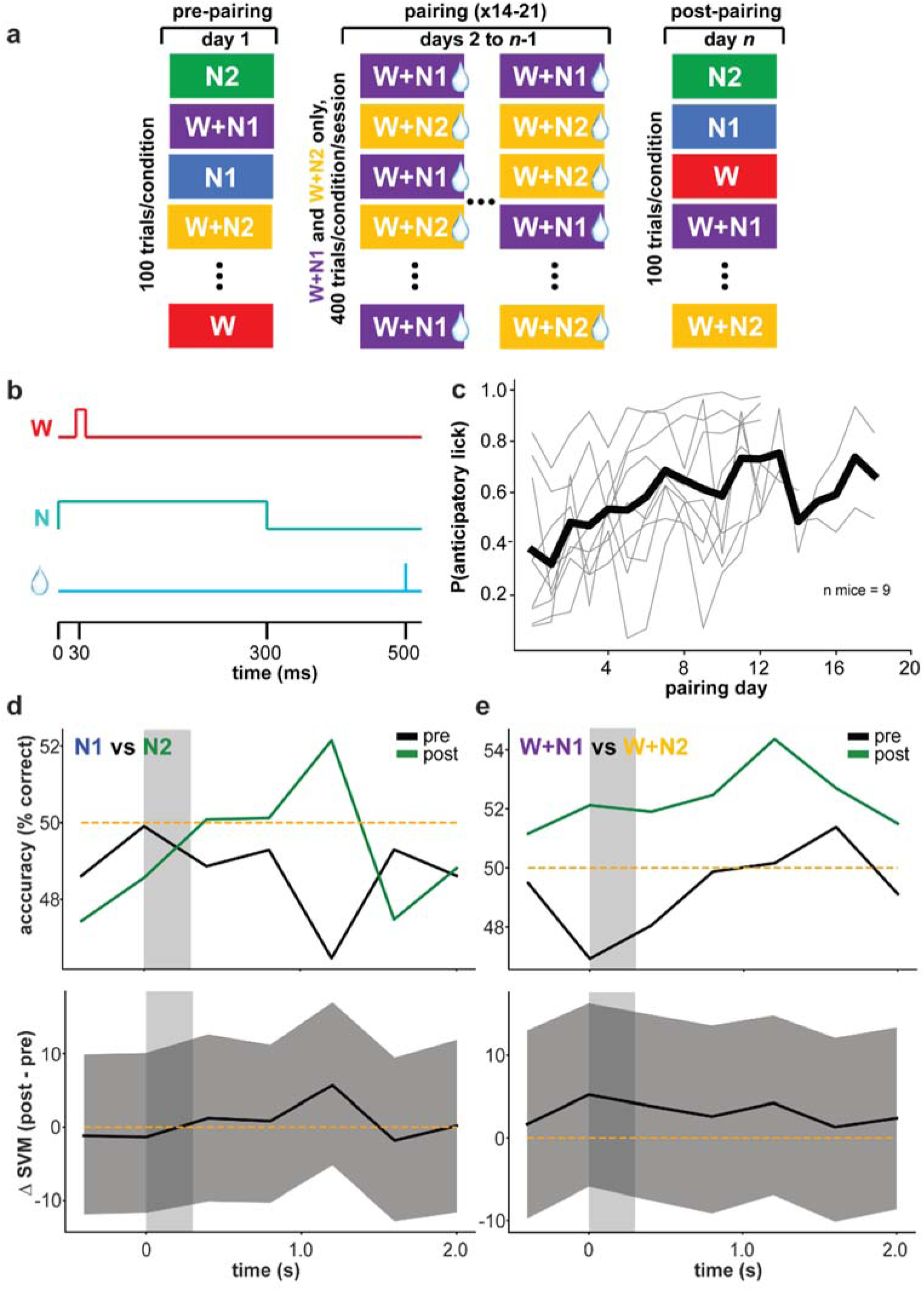
Auditory information in S1 is stable over reinforcement. **a**, Diagram of reward conditioning paradigm. **b**, Diagram of individual trial structure in reward conditioning paradigm. **c**, Probability anticipatory lick vs. pairing day. **d**, Change in N1 vs N2SVM performance after reinforcement. Top panel: mean pre- and post-pairing SVM accuracy over 1000 bootstrapped pre- and post-pairing pseudosession pairs over 9 mice (5,711 cells pre-pairing, 3,079 cells post-pairing), bin size=400 ms. **e** , Same as **d**, but for W+N1vs W+N2.

Mice on average increased anticipatory licking over the course of training (Fig. 7c). We went on to test whether information content about pure auditory stimulus identity changed after reward conditioning by analyzing the change in N1 vs N2 SVM accuracy pre- to post-pairing, finding that the difference was not significantly different from 0 (Fig. 7d; p>0.27, bootstrap test over 1000 iterations). Similarly, W+N1 vs W+N2 SVM performance did change not significantly over the course of reward conditioning (Fig. 7e; p>0.32, bootstrap test over 1000 iterations).For both dichotomies, using a nonlinear MLP rather than a linear decoder yielded similar patterns of results, performing largely around chance levels throughout the trial epoch both before and after reward (Figs. S6, S7). Collectively, these results demonstrate that in stark contrast to whisker inputs (Rabinovich et al. 2022, Benezra et al. under review, Huber et al. 2012), auditory inputs to S1 are remarkably stable in the face of both passive experience and reward conditioning. Thus, while within-modality plasticity is substantial in barrel cortex, cross-modal plasticity as a result of normal experience and learning seems minimal or absent.

## Discussion

In this study, we imaged S1 responses to pure auditory and conjunctive audio-tactile stimuli in both naïve and experienced mice in order to test several hypotheses about the possible functions subserved by cross-modal interactions in the early stages of cortical processing. We first sought to establish whether S1 encodes information about acoustic frequency in naïve mice by measuring neuronal activity evoked by two distinct, intensity-matched, band-limited noise stimuli. Consistent with previous findings (Wallace et al. 2005, Iurilli et al. 2012, Zhang et al. 2020), we observed weak suprathreshold spiking responses to pure auditory stimuli in S1 overall. Moreover, we found that these limited responses encoded minimal information about auditory stimulus identity, despite the fact that the center frequencies for our noise bands are easily discriminable in mice (Koay et al. 2002). Additionally, we found that while presenting acoustic noise concurrent with a tactile stimulus had a significant suppressive effect on whisker contact-evoked responses in S1 overall, the auditory stimuli tested tended to elicit similar patterns of whisker-response modulation across the imaged population, resulting in no significant encoding of specific audio-tactile feature conjunctions.

In order to test whether these stimulus non-specific influences were amenable to modification by different forms of synaptic plasticity, we then subjected mice to various audio-tactile pairing paradigms. Exposing mice to repeated pairings of specific auditory and tactile stimuli presented within tens of milliseconds of each other had no effect on the decodability of either pure auditory or audio-tactile stimulus identity, suggesting that passive experience is insufficient to induce auditory selectivity in S1 even over the course of several days. Furthermore, dispensing reward following presentation of audio-tactile stimulus conjunctions similarly failed to alter the decodability of either pure auditory or audio-tactile stimuli. Notably, this stability in the overall amount of auditory and audio-tactile information encoded by S1 stands in stark contrast to the adaptive, plastic nature of its tactile responses, which show enhanced encoding of rewarded whisker stimuli following reinforcement (Rabinovich et al. 2022, Benezra et al.). These results thus collectively suggest that auditory influences in S1 play a fundamentally non-sensory role, and that they are mediated by qualitatively different circuit and synaptic mechanisms from those subserving the highly plastic, adaptable, stimulus-specific responses to the preferred, tactile sensory modality.

### S1 encodes minimal information about auditory and audio-tactile stimulus identity in naïve mice

Primary sensory cortical areas have long been thought to operate as banks of feature detectors dedicated to extracting granular, low-level perceptual features from a single sensory modality each, but in recent years this view has come under increasing scrutiny. Many studies have shown that even these earliest stages of cortical processing encode a wide variety of variables other than those directly related to unisensory feature detection, including motivational state (Allen et al. 2019, Kauvar & Machado et al. 2020), bodily movement (Musall et al. 2019), task rules (Rodgers & DeWeese 2014, Zempeltzi et al. 2020, Osako 2021), and even spatial location (Saleem et al. 2018). Related work has also shown that primary sensory cortical areas are modified by behavioral relevance and reward contingency (Kato et al. 2015, Poort et al. 2015, Keller et al. 2017, Henschke et al. 2020, Benezra et al. bioRxiv) and reward timing (Shuler et al. 2006, Pantoja et al. 2007, Pleger et al. 2008, Weis et al. 2013, Rabinovich et al. 2022) in particular.

This general reexamination of the role of primary sensory cortical areas has extended to the idea that these regions may be involved in encoding overall sensory context, including integrating information from multiple sensory modalities. Indeed, numerous recent studies have shown that sensory stimuli from non-preferred modalities can modulate activity in every primary sensory neocortical area (Wallace et al. 2005, Higley & Contreras 2005, Banks et al. 2011, Iurilli et al. 2012, Liang et al. 2013, Sieben et al. 2013, Ibrahim et al. 2016, Meijer et al. 2017, Morrill & Hasenstaub 2018, Deneux et al. 2019, Knöpfel et al. 2019, Zhang et al. 2020, Garner et al. 2022, Bimbard et al. 2023). Complementary anatomical and transection studies even suggest that some of these influences may be mediated by direct, monosynaptic connections between primary sensory cortical areas (Budinger et al. 2006, Charbonneau et al. 2012, Iurilli et al. 2012, Stehberg et al. 2014, Henschke et al. 2015). Nevertheless, the functional roles played by these early cross-sensory influences remain enigmatic, and efforts to further elucidate them have yielded contrasting results.

One of the most fundamental constraints on the possible computational roles subserved by these cross-modal, early cortical signals is the extent to which they encode the identity of specific stimuli or combinations thereof. For example, if such signals do not differentially encode distinct stimuli in a region’s non-preferred modality, then they are unlikely to be substantively involved in mapping specific multisensory feature combinations of to appropriate responses, or in predicting future input to one modality based on current input to another modality. We therefore decided to test the hypothesis that sound-evoked activity in S1 encodes information about acoustic frequency, focusing on audio-tactile interactions for their potential ethological relevance; in addition to being extremely important to rodent behavior generally, whisking on different textures has been shown to produce different sounds, thereby opening up the possibility that integrating auditory and tactile information may be useful for disambiguating different materials or denoising degraded vibrissal input (Efron & Lampl 2022, Ernst & Bülthoff 2004, Raposo et al. 2012, Sheppard et al. 2013, Coen et al. bioRxiv). We observed very weak responses to pure auditory stimuli on average, consistent with previous results that auditory input to S1 is mainly inhibitory but with a small, depolarizing late peak (Wallace et al. 2005, Iurilli et al. 2012, Zhang et al. 2020).

Critically, we found that what little activity was observed hardly encoded any information about which of two physiologically and behaviorally intensity-matched acoustic stimuli was presented. These results agree with recent findings by Bimbard et al. (2023), who report that sound-evoked activity in mouse primary visual cortex can be explained almost entirely by uninstructed facial and body movements. Indeed, the present study can be viewed as complementing that work by verifying experimentally that when behavioral differences between auditory stimuli are controlled for, neural responses in non-auditory primary sensory cortical areas largely overlap. Our findings are also broadly consistent with those of Morrill & Hasenstaub (2018), who find that visual grating-evoked responses in primary auditory cortex do not encode variables like stimulus orientation. Collectively, then, these results suggest that apparent cross-modal activity in primary sensory cortical areas chiefly reflects global, stimulus-non-specific signals related to internal state variables like arousal or associated movements. This conclusion contrasts with that of Knöpfel et al. (2019), who find that slightly over a third of sound-responsive cells in primary visual cortex are significantly tuned for acoustic frequency. However, these auditory stimuli were played at a variety of perceived volumes (physical intensity normalized by hearing threshold at that frequency), making it possible that the different V1 responses they elicited were mediated by different behavioral responses as in Bimbard et al. For this reason, our experiments purposefully made use of amplitude-matched auditory stimuli.

Independently of whether S1 encodes pure auditory stimulus identity, we also investigated its capacity to represent different combinations of auditory and tactile features by testing whether different concurrently presented sound-and-whisker stimulus pairs could be decoded from S1 activity. Theoretical work has repeatedly shown that such nonlinear mixed selectivity is critical for solving complex tasks (Rigotti et al. 2013, Fusi et al. 2016), and nonlinear mixing is known to occur within sensory modalities in primary sensory cortical areas (Lavzin et al. 2012, Xu et al. 2012, Nogueira et al. 2023). Moreover, one might expect performance on specifically multisensory complex tasks to benefit from mixing cross-sensory inputs in areas where receptive fields are small and sensory information is represented in granular detail, like primary sensory cortical areas. Consistent with Ibrahim et al. (2016) and Zhang et al. (2020), we found that playing sounds concurrent with whisker stimulation had a significant net inhibitory effect on whisker-evoked responses in S1. We extend that work, however, by showing that rather than selectively suppressing whisker responses in distinct subpopulations of S1, noise bursts of different frequency ranges tend to suppress the same populations of cells, largely precluding a combinatorial code for specific audio-tactile stimuli in S1. Thus, our initial experiments revealed very little information about the identity of either pure auditory or audio-tactile stimuli encoded in S1.

These results appear to contrast somewhat with those of Deneux et al. (2019), who report both that different auditory stimuli evoke distinct population-averaged responses in primary visual cortex and that these sound-evoked responses depend on concurrent visual input. However, these stimuli differed in their intensity profiles over time, with one abruptly beginning loudly and gradually ramping down in volume over time while the other began faintly and gradually intensified, making it possible that the different responses to these stimuli were mediated by different behavioral (e.g. startle or arousal) responses. Indeed, our results along with those of Bimbard et al. suggest that when such behavioral effects are controlled for, little information about cross-modal stimuli *per se* remains in primary sensory cortical areas. Additionally, Renard et al. (2022) suggest that odorant identity can be decoded from barrel cortex even in the absence of reafferent whisking activity following facial nerve transection. Nonetheless, these results could be at least partly accounted for by other, non-facial bodily movements (Musall et al. 2018, Kauvar and Machado et al. 2020, Bimbard et al. 2023), efference copy, or intrinsic affective valence odor. On the other hand, this contrast may reflect genuine differences between audio-tactile and olfactory-tactile processing, especially since the first stage of cortical olfactory information processing is the piriform, part of the phylogenetically older and perhaps qualitatively distinct paleocortical region of the cortical mantle (Klingler 2017).

The suppressive effect of auditory stimuli on tactile responses that we and Zhang et al. (2020) describe contrast with those of Godenzini et al. (2021), who observed that auditory stimuli had a net facilitative effect on forepaw responses in S1. These differences may be due to their measuring responses in the forepaw region of S1 rather than the barrel field or applying repetitive tactile stimuli as well as noise burst stimulus with a much wider passband. That study used only a single auditory stimulus and did not examine the decodability of auditory stimuli.

### Auditory information content of sound-evoked signals in S1 is stable over experience

After finding that sound-evoked activity in S1 encodes little stimulus-specific information in naïve mice, we considered the possibility that these signals are sensitive to the statistics of the multisensory environment and selectively reflect correlations between auditory and tactile inputs learned through experience. We therefore went on to investigate whether the information-coding properties of these weak, generic inputs were amenable to modification via experience. We first tested the hypothesis that merely pairing a given auditory stimulus with a tactile stimulus in a passive setting would suffice to engage Hebbian-like plasticity and result in the paired auditory stimulus specifically reactivating the same ensemble of neurons driven by the tactile stimulus. Indeed, previous research has shown that such pattern completion occurs between stimuli of the same sensory modality in primary sensory cortical areas (Xu et al. 2012, Gavornik & Bear 2014, Carillo-Reid et al. 2016, Libby & Buschman 2021), and has long been hypothesized to occur across sensory modalities as well (Durup & Fessard 1935, Vaudano et al. 1990). However, we found little evidence of any such mechanism at play between auditory and tactile inputs to S1, even after presenting over one thousand pairings and spacing them out over several days to allow for the consolidative effects of sleep (Grewe et al. 2017).

We also considered the hypothesis that passive exposure to correlations between specific sounds and whisker stimulation might induce nonlinear mixing otherwise absent in naïve mice, giving rise to enhanced representations of commonly encountered combinations of auditory and tactile stimuli. Indeed, theoretical work suggests that enriched selectivity for commonly occurring stimuli may improve neural coding efficiency (Deep & Simoncelli, bioRxiv), and physiological studies have shown that dendritic nonlinearities can be induced through repeated coincident stimulation of pre- and postsynaptic cells in other cortical structures like hippocampus (Brandalise et al. 2016). However, when we trained a linear decoder to classify different audio-tactile stimulus conjunctions after pairing, performance was not significantly improved compared to baseline, suggesting that passive experience was insufficient to induce such specialized multisensory representations in S1.

Once again these results contrast with those of Knöpfel et al., who report that passively pairing an auditory with a visual stimulus specifically increases V1 responses to the paired auditory stimulus. However, because that study focused on mean responses rather than patterns of population activity, it remains possible that the observed increase was due to a generic orienting or attentional response rather than to Hebbian-like plasticity leading the auditory stimulus to reactivate the ensemble representing the paired visual stimulus. Alternatively, differences between that work and the present study may be reflective of genuine differences between primary visual and somatosensory cortical areas in mouse.

Finally, we tested the hypothesis that reinforcement with reward may be necessary to induce plasticity and enhance encoding of auditory stimulus features in S1. Indeed, for stimuli *within* a primary sensory cortical area’s preferred modality, reward-pairing has repeatedly been shown to strongly and robustly enhance encoding of rewarded stimuli (Kato et al. 2015, Poort et al. 2015, Keller et al. 2017, Lacefield et al. 2019, Henschke et al. 2020, Benezra et al. bioRxiv) and even of related features like expected reward timing (Shuler et al. 2006, Pantoja et al. 2007, Pleger et al. 2008, Weis et al. 2013, Rabinovich et al. 2022). However, we found by contrast that in the multisensory case, linear decoder performance for both pure auditory and conjunctive audio-tactile stimuli in S1 remained near chance levels even after up to three weeks of following conjunctive audio-tactile stimulus pairs with reward.

Garner et al. (2022) found that when a particular auditory-visual stimulus sequence predicts reward, V1 responses to the visual stimulus are selectively suppressed by the predictive auditory stimulus. In that study, however, only one auditory stimulus predicted reward and thus elicited licking, making it possible that the selective suppression of visual responses by the reward-predictive sound was mediated by enhanced consummatory behavior (though suppression wasn’t observed on false alarm trials, higher lick rates on hit trials may have nevertheless been partly responsible for the effect). In our study, by contrast, reinforcement with equal reward contingencies across stimuli failed to result in enhanced auditory encoding in S1, despite the fact that equally rewarding two stimuli within a primary sensory cortical area’s preferred modality has been shown to result in improved decoder performance (Henschke et al. 2020). Alternatively, these contrasting results may reflect genuine differences between audio-visual and audio-tactile processing; indeed, anatomical studies have shown anisotropies in the number of projections in the A1-V1 and A1-S1 pathways in other rodent species (Henschke et al. 2015). Finally, it is worth noting that all pairing experiments in the present study employed the same latency between auditory and tactile stimuli, and that altering this latency could affect multisensory integration, similar to how Meijer et al. (2017) found that temporal congruency between amplitude-modulated auditory and visual stimuli affected the degree of sound-driven modulation of visual responses.

Despite encoding little to no stimulus-specific information *per se*, cross-sensory influences in primary sensory cortical areas may nonetheless play an important role in shaping early sensory responses. For example, thalamocortical synapses mediating input from a primary sensory cortical area’s preferred modality may control the distribution of that area’s receptive field centers by undergoing relatively long-lasting changes that privilege representation of behaviorally relevant sensory features, while corticocortical synapses conveying information about movement, internal state, or the general presence of input from another modality may control the width of those receptive fields via short-term, instantly reversible subtractive or divisive gain changes (Higley & Contreras 2005, Priebe & Fester 2006, Iurilli et al. 2012, Schneider et al. 2014, Zhou et al. 2014, Zhang et al. 2020). Theoretical work suggests that even such response sharpening and normalization can subserve selective attention and improve stimulus encoding when inputs are low-dimensional (e.g. when inputs from different modalities are temporally correlated or coincident; Zhang & Sejnowski 1999, Brown & Bäcker 2006, Carandini & Heeger 2012), and experimental work has indeed shown that this type of suppression can improve stimulus discriminability and even performance on behavioral tasks (Brenner et al. 2000, Kok et al. 2012, Zhou et al. 2014). Thus, while primary sensory cortical areas may in the end chiefly encode sensory input from a single preferred modality as long held, their activity may yet be indirectly albeit importantly shaped by other modalities as well.

Overall then, our results suggest that sound-evoked activity S1 is largely stimulus non-specific, consistent with these responses reflecting movement-related or internal state variables rather than the sensory qualities of auditory stimuli *per se*. Furthermore, we find that the information content of these inputs is extremely stable in the face of various forms of learning and experience. This latter finding contrasts starkly with previous studies showing that representations of a primary sensory cortical area’s preferred modality are remarkably plastic, adapting dynamically to task demands and the statistics of the sensory environment (Shuler et al. 2006, Pantoja et al. 2007, Pleger et al. 2008, Xu et al. 2012, Weis et al. 2013, Gavornik & Bear 2014, Martins & Froemke 2015, Kato et al. 2015, Poort et al. 2015, Keller et al. 2017, Lacefield et al. 2019, Henschke et al. 2020, Rabinovich et al. 2022, Benezra et al. bioRxiv). This remarkable disparity between the adaptability of inputs related to a primary cortical area’s preferred and non-preferred sensory modalities suggests that these inputs may be mediated by qualitatively different types of synapses marked by distinct types of receptors, ion channels, dendritic transients, and plasticity rules subserving different functional and computational roles.

## Materials and methods

### Subjects

All experiments were approved by the Columbia University Institutional Animal Care and Use Committee. Subjects consisted of 13 wild-type CBA/J mice housed in groups of up to 5 littermates per cage. Mice had *ad libitum* access to food throughout the course of all experiments, and running wheels were placed in home cages for enrichment. Any mice observed barbering cage mates were separated and singly housed. During pairing, mice in the reinforcement condition were water-restricted and weighed daily to ensure adequate weight was maintained; if body weight fell to less than 80% of pre-training baseline level, then mice were given 5 additional minutes of free access to water in their home cage.

### Virus injection and cranial window implant

Mice were intracranially injected in the left primary somatosensory cortex with adeno-associated virus encoding the genetically-encoded calcium indicator GCaMP6f. Twelve out of 13 mice were injected with GCaMP6f under the control of the Synapsin promoter (pENN.AAV9.Syn.GCamP6f.WPRE.SV40, nominal titer 2.80 x 10^13^ gc/mL), while the remaining mouse was injected with GCaMP6f under the control the CaMKII promoter (pENN.AAV5.CamKII.GCaMP6f.WPRE.SV40, nominal titer 2.30 x 10^13^ gc/mL). During all surgeries, anesthesia was induced using 3% isoflurane then maintained with 1-2% isoflurane while body temperature was maintained at ∼37.0° C using a homeothermic blanket. Once anesthetized, mice were placed in a stereotax, and eyes were kept lubricated with ophthalmic ointment. Toe pinch was used to assess depth of anesthesia every 15 minutes, and the concentration of isoflurane was titrated to eliminate the hindpaw withdrawal reflex.

In mice injected with GCamP6f under the control of the Synapsin promoter, virus injection and cranial window implantation were performed in the same session. Mice were injected intramuscularly with 2 mg/kg dexamethasone 3-4 hours prior to surgery, then anesthetized and administered subcutaneous 0.1 mg/kg buprenorphine and 6 mg/kg bupivacaine as analgesia. The entire scalp was shaved and removed, and a ∼4-mm diameter craniotomy was made in the skull centered around 1.5 mm posterior to bregma and 3.5 mm lateral of the midline. A pipette was then inserted ∼250 µm into the brain at 2 or 3 different sites spanning the area of the craniotomy. Approximately 50 nL of virus solution was injected at each location over 3 pulses spaced one minute apart (for a total of 150 nL at each location?). After the last pulse, the pipette was left in the brain for 3 minutes before being withdrawn. A 4-mm diameter glass coverslip was placed over the craniotomy and affixed in place with cyanoacrylate, and a headpost was bonded to the skull with dental cement. Following surgery, mice were subcutaneously administered buprenorphine approximately every 12 hours for 48 hours and weighed every day for 5 days. If body weight fell below 80% of pre-surgical levels, food would be dampened and placed on the home cage floor for easy access and supplemented with hydrogel. If weight loss persisted, mice were euthanized.

In mice injected with GCaMP6f under the control of the CaMKII promoter, virus injection and cranial window implantation were performed during separate surgery sessions. In addition to the anesthesia and analgesia described above, mice were administered 5 mg/kg subcutaneous carprofen prior to surgery. A small incision was made in the scalp instead of removing it entirely, a small region of skull was thinned in a single location over the center of the barrel field. A pipette was inserted through the thinned skull and 150 nL of virus solution were injected at 150 and 300 µm below the pial surface. At each depth, injections were delivered in 3 equal pulses with one minute between pulses, and after each sequence of 3 pulses, the pipette was left in place for 3 minutes before being withdrawn. The thinned region of skull was then covered with a bolus of artificial cerebrospinal fluid and sealed with cyanoacrylate, and the incision sutured and glued shut using vetbond. Mice were given subcutaneous carprofen 24 hours later, then the virus was then allowed to express for approximately 3 months. Following this incubation period, cranial window and headpost implantation proceeded as described previously.

### Intrinsic signal optical imaging

After cranial window implantation, the location of barrel cortex was confirmed using intrinsic signal optical imaging, during which the red-wavelength reflectance of the brain was imaged while whiskers were stimulated with a piezoelectric cantilever. Mice were anesthetized with isoflurane, administered ophthalmic ointment, and body temperature was maintained at approximately 37.0° C. The brain was illuminated with a 700-nm incandescent light, and whiskers were stimulated one at a time in 5 Hz pulses with a 10-second interstimulus interval between trials of 10 pulses each for ∼5-20 trials depending on how many trials were necessary to observe discernible barrel structures. Images were acquired at a spatial resolution of 512 x 512 pixels using a 5x/0.16 NA objective and Rolera MGi Plus digital camera. Images acquired during whisker stimulation were subtracted from a mean baseline image and averaged over trials to generate maps of changes in cortical reflectance evoked by deflection of each whisker, indicating the location of the corresponding barrel.

### 2-photon calcium imaging

Once the location of the barrel field had been identified, mice were head fixed and imaged with tunable laser set to 940 nm through a 16x/0.8NA water immersion Nikon objective. The beam was swept by a resonant scanner at a frequency of approximately 30 frames per second at 2x digital zoom over an effective field of view of approximately 480 x 480 μm, and data were collected using the ScanImage software package at a spatial resolution of 512 x 512 pixels. For each mouse, a suitable 2-photon imaging site was selected based on location within the barrel field, number of cells, quality of expression, and relative absence of occluding vasculature. Most imaging sites were in the C or D row as assessed by intrinsic signal optical imaging.

For each mouse, the same cells were located at the beginning of each session using vascular and cellular landmarks and imaged over the course of the entire pairing paradigm. All mice were imaged during pre- and post-pairing probe sessions as well on the first and last pairing sessions. Additionally, mice in the unrewarded condition were imaged during every pairing session. Mice in the rewarded condition were imaged on alternate pairing days in order to reduce the possibility of thermal tissue damage and phototoxicity.

### Stimulus pairing paradigm

Mice were head-fixed under the 2-photon microscope and presented with trials consisting of either a tactile stimulus, an auditory stimulus, or a conjunctive audio-tactile stimulus. Tactile stimuli consisted of a pole moved through the whisker field by a stepper motor that rotated a full 360°. The motor was under the control of a SilentStepStick TMC 2100 stepper driver designed to minimize acoustic noise, deflecting the whiskers at an angular velocity of about 1800° per second. Auditory stimuli consisted of 300 ms band-limited noise bursts of one of two frequency ranges: a lower-frequency range from 8.5-10.5 kHz, and a higher-frequency range from 16.5-18.5 kHz. Auditory stimuli were generated randomly on each trial by a data acquisition board (National Instruments) with an output sample rate of 100 kHz and played through an 0.25W, 8Ω speaker at a volume of approximately 70 dB SPL_A_ positioned ∼4.5 cm from the mouse’s ear. Conjunctive audio-tactile stimuli consisted of an auditory stimulus followed by a tactile stimulus contacting the whiskers approximately 30 ms later, with some variability due to whisker movement. In some additional trials neither sound was presented.

Trials were presented with a minimum inter-trial interval of 1.2 seconds plus an additional interval drawn from an exponential distribution with a mean of 0.3 seconds, to prevent mice from predicting the time of stimulus presentation. In order to prevent stepper noise or vibration from becoming predictive of the whisker stimulus and potentially “blocking” any association between auditory and tactile stimuli, the stepper was moved on all trials. On trials with no whisker stimulus (sound-only and no-stimulus trials), the pole was moved away from the mouse’s face. In total, 6 different stimulus conditions were presented: whisker stimulus alone (W), auditory stimulus 1 alone (N1), sound 2 alone (N2), whisker stimulus with sound 1 (W+N1), whisker stimulus with sound 2 (W+N2), and no stimulus (NS).

Mice were habituated to head fixation for a week before imaging. During habituation sessions, the scanner was run with the objective in place to acclimate mice to the ambient noise of the 2-photon setup as well. Mice were then imaged during a pre-pairing probe session consisting of randomly interleaved trials of all stimulus conditions (100 trials/condition) the day before pairing began. During the pairing phase, mice in the unrewarded paradigm received 3 consecutive days of pairing sessions, each of which consisted of 800 randomly interleaved trials of W+N1and N2 (400 trials/condition). After pairing, mice were presented with an additional post-pairing probe session of randomly interleaved trials from all conditions.

In the rewarded paradigm, mice were water-restricted for a week before pairing. In addition to habituation to head-fixation, behavior was shaped by delivering water from a lick port every time the mouse licked. After an initial pre-pairing session consisting of unrewarded stimuli of all conditions, mice received daily pairing sessions consisting of randomly interleaved W+N1and W+N2 trials (400 trials/condition). In order to ensure that any changes in decoder performance were due to changes in the sensory representation of the stimuli *per se* rather than to changes in behavior or arousal state, trials of both conditions were followed by a water reward after a 200-ms trace interval. Licking was measured using a capacitive touch sensor connected to the lick port, and anticipatory licking during pairing sessions was quantified as the probability of at least one lick occurring during the trace interval. Mice received pairing sessions every day for at least 2 weeks. If the probability of anticipatory licking was at least 0.7 for 2 or more consecutive sessions at the end of 2 weeks, pairing was halted and mice were given a post-pairing probe session consisting of random trials of all conditions with no reward. If the probability of anticipatory licking was not at least 0.7 for the previous 2 days at the end of 2 weeks, then mice were given an additional week of pairing sessions, after which they were given a post-pairing probe session.

### Auditory brainstem response recordings

To verify that both auditory stimuli evoked similar responses at the level of the cochlea and brainstem, auditory brainstem response recordings were performed on each mouse after pairing was complete. Mice were anesthetized with 70 mg/kg pentobarbital and body temperature was maintained at ∼37.0° C. Because mice had been implanted with headplates and cranial windows, electrodes could not be implanted under the scalp as in typical auditory brainstem response recordings, so the skull was thinned instead to allow for a silver wire electrode to be implanted directly into the brain. A reference electrode was inserted at the base of the left pinna, and the tail was grounded. Signals were amplified with a 20x gain battery-powered preamp (Thomas Recording) and then with a Lynx amplifier set to 3004x gain with low and high cutoff frequencies of 300 and 3000 Hz, respectively. Stimuli consisted of the same band-limited noise bursts used during imaging, but shortened from 300 to 100 ms to allow for more trials per recording session. Each stimulus was presented 1000 times, and data for each condition were averaged across trials.

### Quantifying cellular activity

Raw 2-photon imaging data were motion corrected and segmented into individual cells using the Suite2p software package (Pachitariu et al. 2017), and data on each trial were ΔF/F normalized to mean fluorescence over 0.5 seconds before stimulus onset. Additionally, ΔF/F time series were smoothed using a rolling boxcar filter averaging each frame with the previous 2-3 frames. Cells were classified as significantly responsive to a stimulus if peak ΔF/F over the 1.25 s following stimulus onset was significantly different from peak ΔF/F over the 0.75 s immediately preceding stimulus onset in a rank sum test. In order to assess whether significant transients occurred on single trials, a rank sum test was performed on the distribution of ΔF/F values from 0.75 s immediately preceding stimulus onset versus 1.25 s immediately following stimulus onset on the same trial. For analyses requiring separate tests for each neuron, including calculating the percentage of cells significantly responsive to different stimuli and the percentage of W-responsive cells significantly modulated by N1 and N2, all p-values were adjusted for false discovery rate using the Benjamini-Hochberg procedure.

### Support vector machine analyses

For support vector machine (SVM) analyses, data were trialized and binned into 300-400 ms time bins. SVMs were trained and tested using the Python sklearn.svm.LinearSVC module with a variety of different hyperparameters, including bin size, penalty (L1 or L2), regularization strength, and prestimulus period used to compute ΔF/F. Different hyperparameters worked best for different analyses, so hyperparameters were searched separately for each analysis and selected to maximize SVM accuracy.

To circumvent overfitting problems caused by the fact that most sessions included data from more cells than trials, data from different mice were randomly subsampled with replacement and concatenated into pseudosimultaneous trials (“pseudotrials”). This entailed first splitting trials by condition then randomly partitioning them into non-overlapping subsets of training and test trials for each mouse, with an 80/20 split between training and test. Training pseudotrials of a given condition were then generated by randomly sampling (with replacement) one training trial of that condition from each mouse and concatenating the neural activity from those trials into a *ν*-by-*b* matrix, where *ν* is the total number of cells across mice and *b* is the number of time bins. This procedure was repeated 200 times per condition for both training and test to yield a single “pseudosession.” Training and testing an SVM on a single pseudosession yielded one accuracy value per time bin, and the whole procedure was repeated over 50 pseudosessions and accuracy averaged over repetitions to obtain a robust estimate of SVM performance at each time bin.

In order to test whether SVM performance on pseudosession data was significantly above chance, trial labels were shuffled, and each shuffle was used to generate 50 pseudosessions using the same procedure described above. Accuracy was averaged across pseudosessions to yield a single estimate of SVM performance per shuffle per time bin. This procedure was iterated 1000 times to yield a shuffle distribution for each time bin. SVM performance was deemed significantly above chance at a given time bin if performance was above the 95th percentile of the shuffle distribution.

To test whether SVM accuracy changed significantly over the course of pairing, 1000 pairs of pseudosessions were generated, one from pre-pairing data and one from post-pairing data, and post-pairing accuracy minus pre-paring accuracy was computed for each time bin. This resulted in a bootstrap distribution of differences, and SVM performance was said to have increased significantly at a given time bin if 0 was below the 5^th^ percentile of the bootstrap distribution for that time bin.

For multilayer perceptron (MLP) analyses, data were binned, randomly sampled with replacement, and concatenated across mice into pseudosimultaneous sessions as for the SVMs. MLPs were trained and tested using the sklearn.neural_network.MLPClassifier module. Results for all classifications were qualitatively similar across a wide variety of network architectures and regularization hyperparameters, so all plots of MLP performance show results for a network with two hidden layers of 50 units each and an L2 regularization strength of 10^-3^. Since mean MLP performance across pseudosessions was always within approximately three percentage points of chance performance for all comparisons pre- and post-pairing, and because training and testing an MLP can take orders of magnitude longer than training and testing an SVM for the same comparison, bootstrap tests were forgone in favor of simply plotting the standard deviation of MLP performance across pseudosessions. One of nine mice was omitted from the post-reward pairing MLPs due to the low number of trials in the final probe session (approximately 1/3 of that in all other mice) along with the sensitivity of the nonlinear decoder to random clustering of certain trial types towards the beginning of the session, when fluorescence signals may differ from those during the rest of the session as a result of photobleaching or movement related to acclimation to head-fixation.

### Whisking analysis

Video of the whiskers was recorded during all imaging and pairing sessions with a PS3Eye webcam at a frame rate of 125 frames per second. Whiskers were automatically segmented and tracked using the Whisk software package (Clack et al. 2012) and manually curated using custom software. Median whisker angle was computed for every frame then used to compute trial-averaged whisker position time series for each condition and each mouse.

## ACKNOWLEDGMENTS

The authors thank Rui Ponte Costa and Kerry Walker for comments on the manuscript. Support was provided by a Wellcome Trust Discovery Award, an Academy of Medical Sciences Professorship, NIH/NINDS R01 NS069679 and R01 NS094659 (RMB); and NIH/NINDS F31 NS105490 and the Kavli Institute for Brain Science (DDK).

---

**Figure S1:**
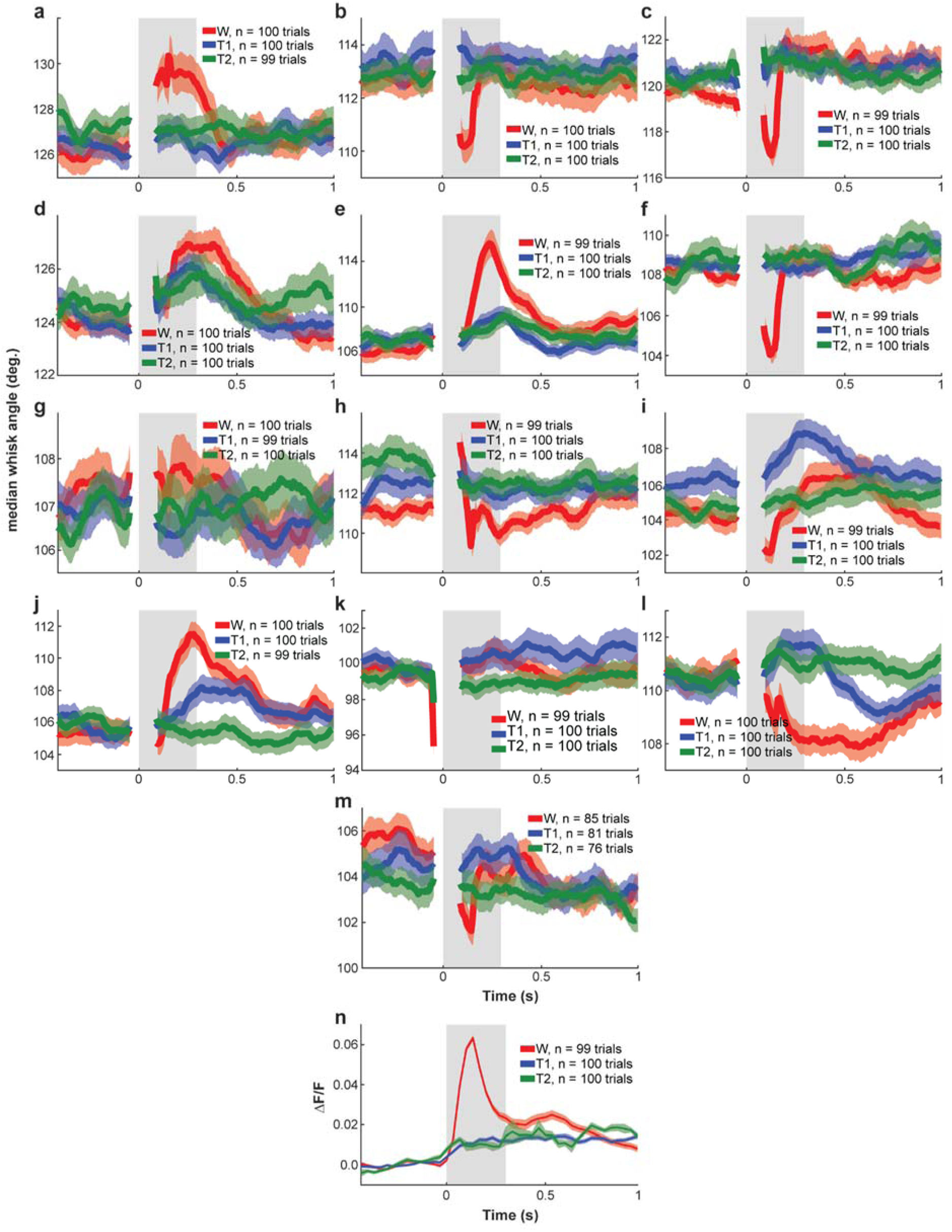
Trial-averaged whisking responses to auditory stimuli N1 and N2 are highly similar. **a-m,** Trial-averaged median whisker angle vs time in response to W, N1, and N2 during initial imaging session for all 13 individual mice used in the study. Each plot represents one mouse. Measurements at each time point represent median angle across all whiskers relative to the mouse’s whisker pad within the video image plane, in degrees. Red: whisker alone, blue: N1, green: N2. Bold traces: trial-averaged median whisker angle; shaded region: standard error of the mean. **n,** Population-averaged neural response to W, N1, and N2 for mouse 3285-3 (whisking depicted in panel **i**).

**Figure S2:**
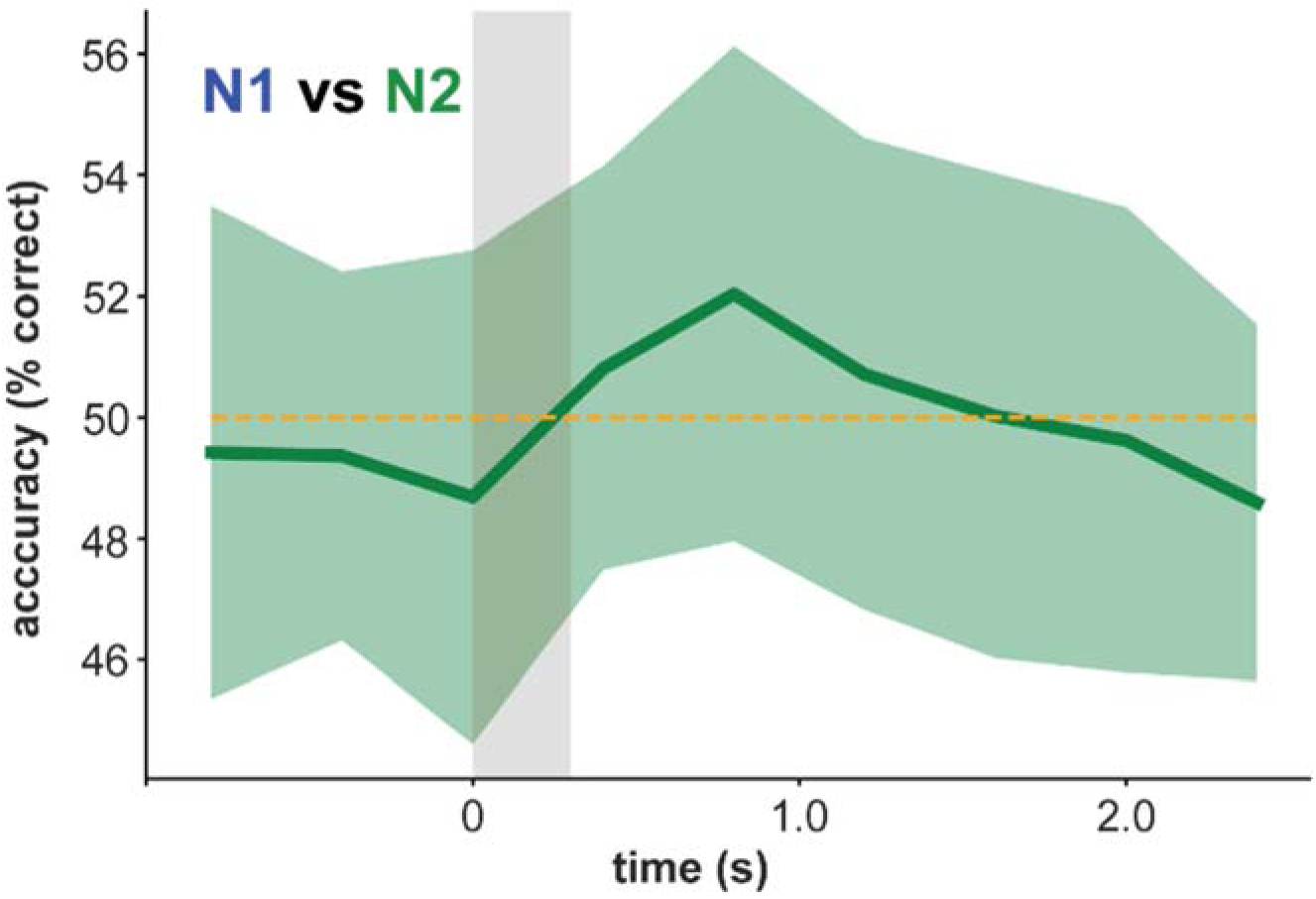
N1 vs N2 multilayer perceptron trained on initial imaging session data performs around chance. Cross-validated N1 vs N2 multilayer perceptron (MLP) accuracy vs. time for initial imaging session. Bin size equals 400 ms. Bold line: mean cross-validated MLP accuracy across 100 pseudosimultaneous sessions generated from 13 mice and 7,376 neurons plus associated neuropil regions-of-interest for each (see methods). Shaded error region: standard deviation across 100 pseudosessions. Shaded rectangle: auditory stimulus epoch. Yellow dashed line: chance performance level.

**Figure S3:**
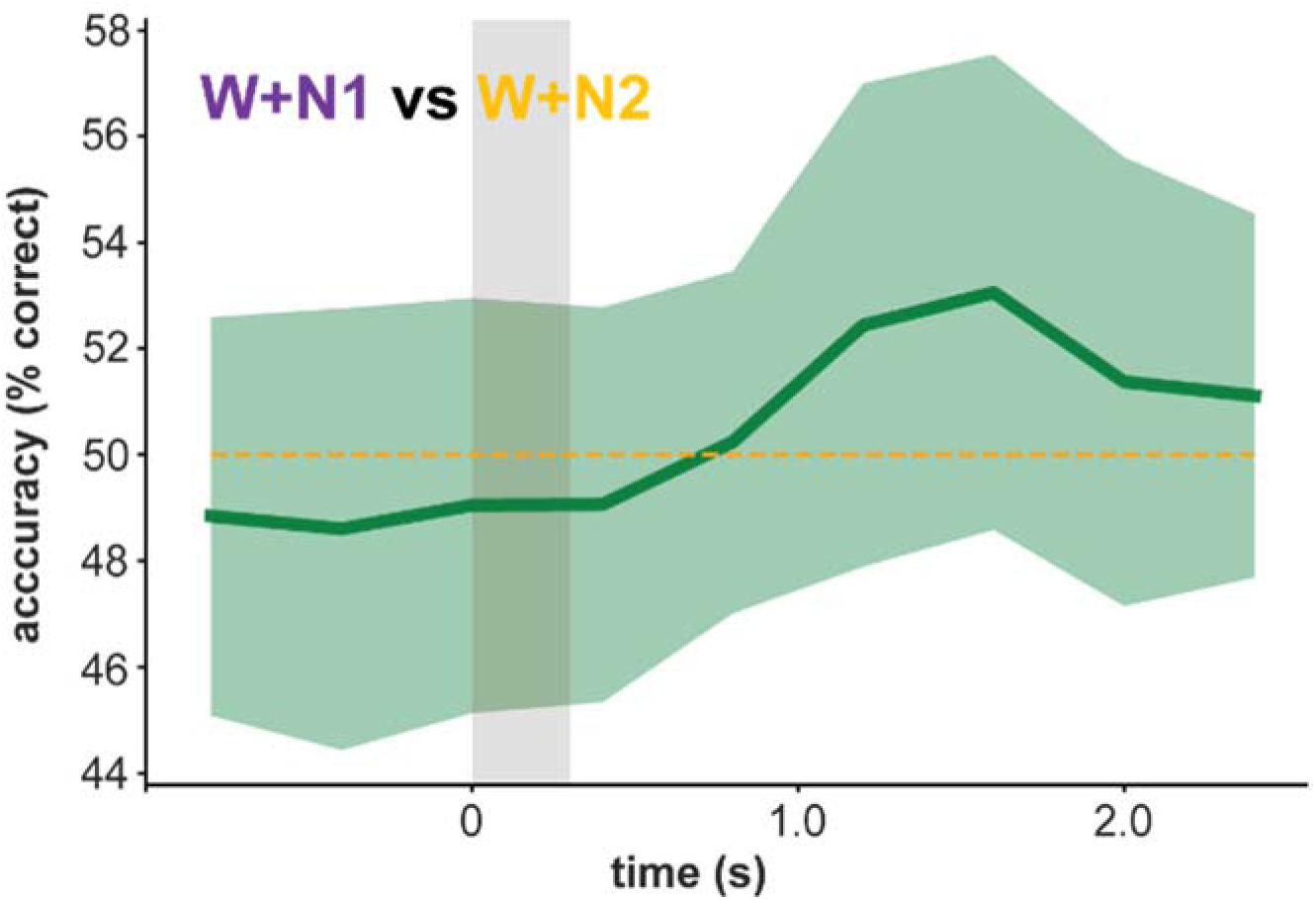
W+N1 vs W+N2 multilayer perceptron trained on initial imaging session data performs around chance. Cross-validated W+N1 vs W+N2 MLP accuracy vs. time for initial imaging session. Bin size equals 400 ms. Bold line: mean cross-validated MLP accuracy across 100 pseudosimultaneous sessions generated from 13 mice and 7,376 neurons plus associated neuropil regions-of-interest for each (see methods). Shaded error region: standard deviation across 100 pseudosessions. Shaded rectangle: auditory stimulus epoch. Yellow dashed line: chance performance level.

**Figure S4:**
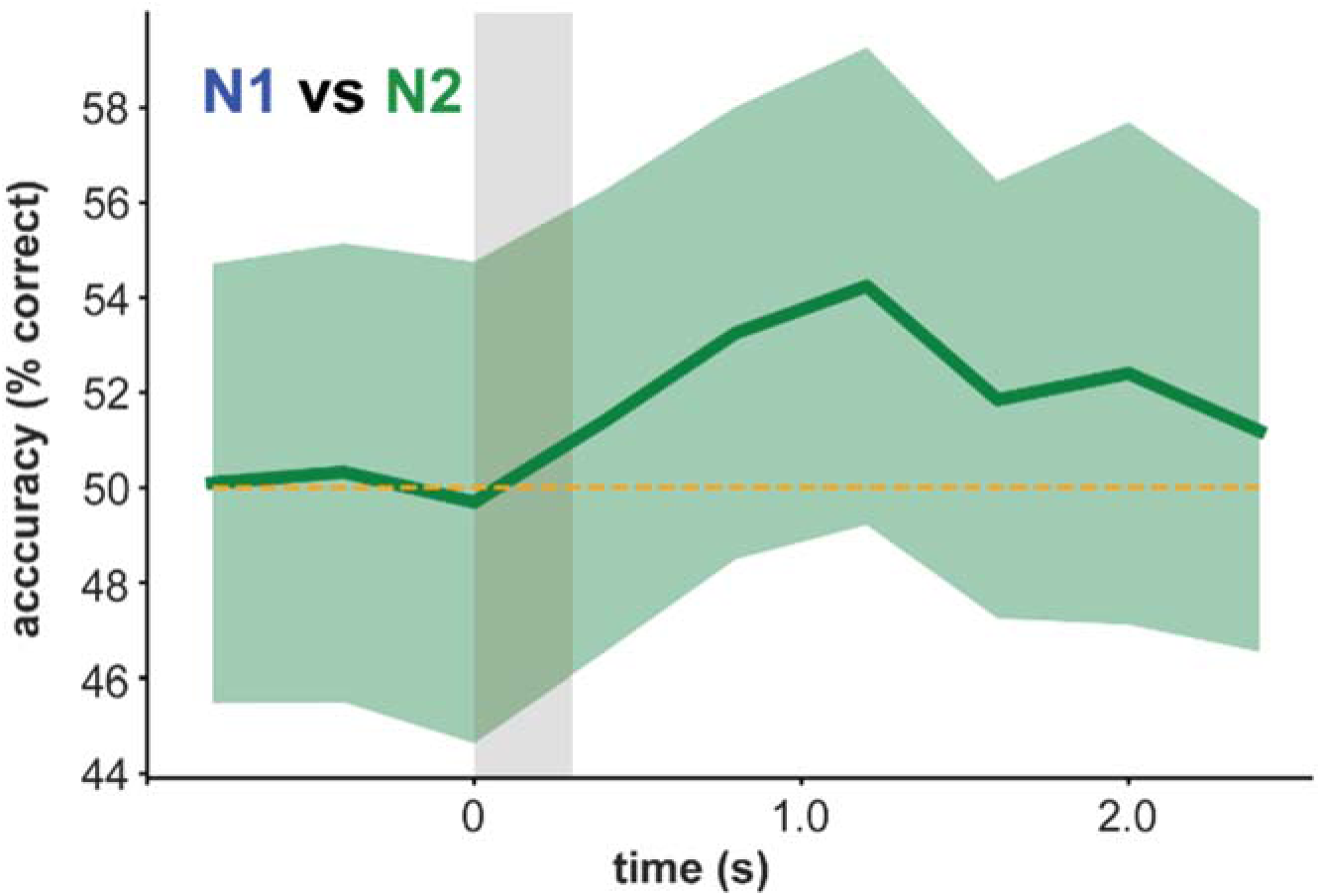
N1 vs N2 multilayer perceptron performs around chance after passive pairing. Cross-validated N1 vs N2 MLP accuracy vs. time after passive pairing of W and N1. Bold line: mean cross-validated MLP accuracy across 100 pseudosimultaneous sessions generated from 4 mice and 1,184 neurons plus associated neuropil regions-of-interest for each (see methods). Shaded error region: standard deviation across 100 pseudosessions. Shaded rectangle: auditory stimulus epoch. Yellow dashed line: chance performance level.

**Figure S5:**
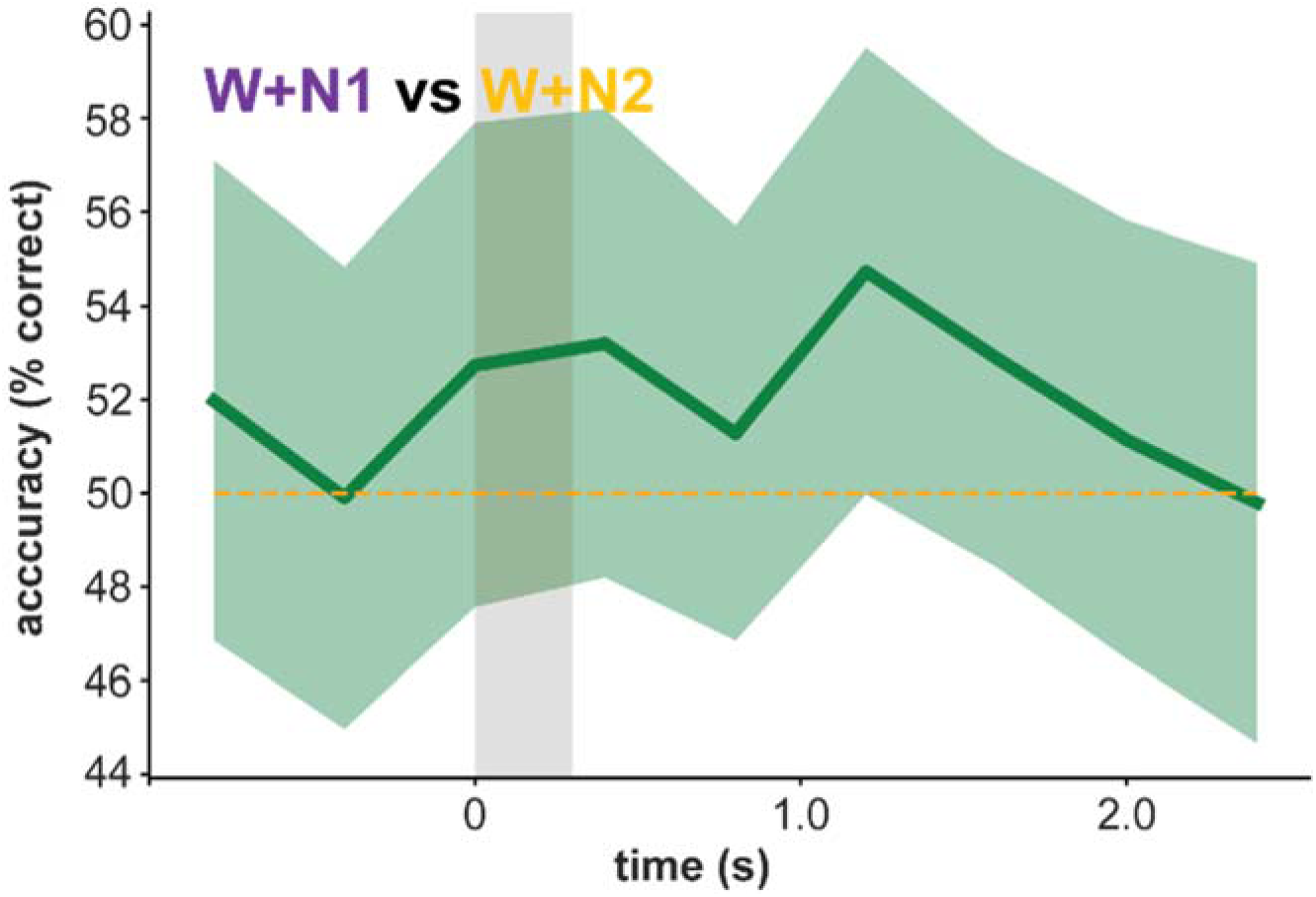
W+N1 vs W+N2 multilayer perceptron performs around chance after passive pairing. Cross-validated W+N1 vs W+N2 MLP accuracy vs. time after passive pairing of W and N1. Bin size equals 400 ms. Bold line: mean cross-validated MLP accuracy across 100 pseudosimultaneous sessions generated from 4 mice and 1,184 neurons plus associated neuropil regions-of-interest for each (see methods). Shaded error region: standard deviation across 100 pseudosessions. Shaded rectangle: auditory stimulus epoch. Yellow dashed line: chance performance level.

**Figure S6:**
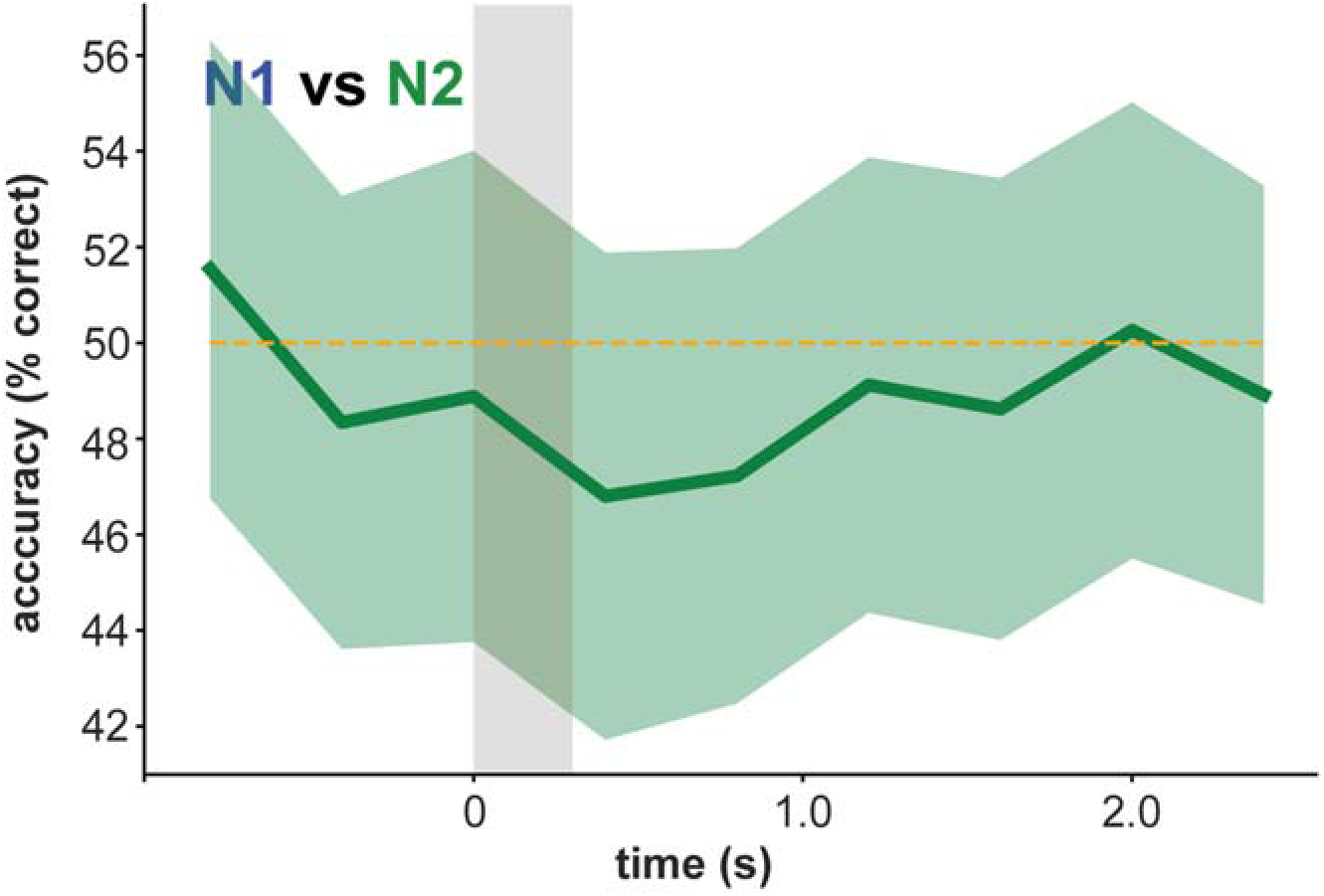
N1 vs N2 multilayer perceptron performs around chance after reward pairing. Cross-validated N1 vs N2 MLP accuracy vs. time after pairing both W+N1 and W+N2 with reward. Bin size equals 400 ms. Bold line: mean cross-validated MLP accuracy across 100 pseudosimultaneous sessions generated from 8 mice and 2,926 neurons plus associated neuropil regions-of-interest for each (see methods). Shaded error region: standard deviation across 100 pseudosessions. Shaded rectangle: auditory stimulus epoch. Yellow dashed line: chance performance level.

**Figure S7:**
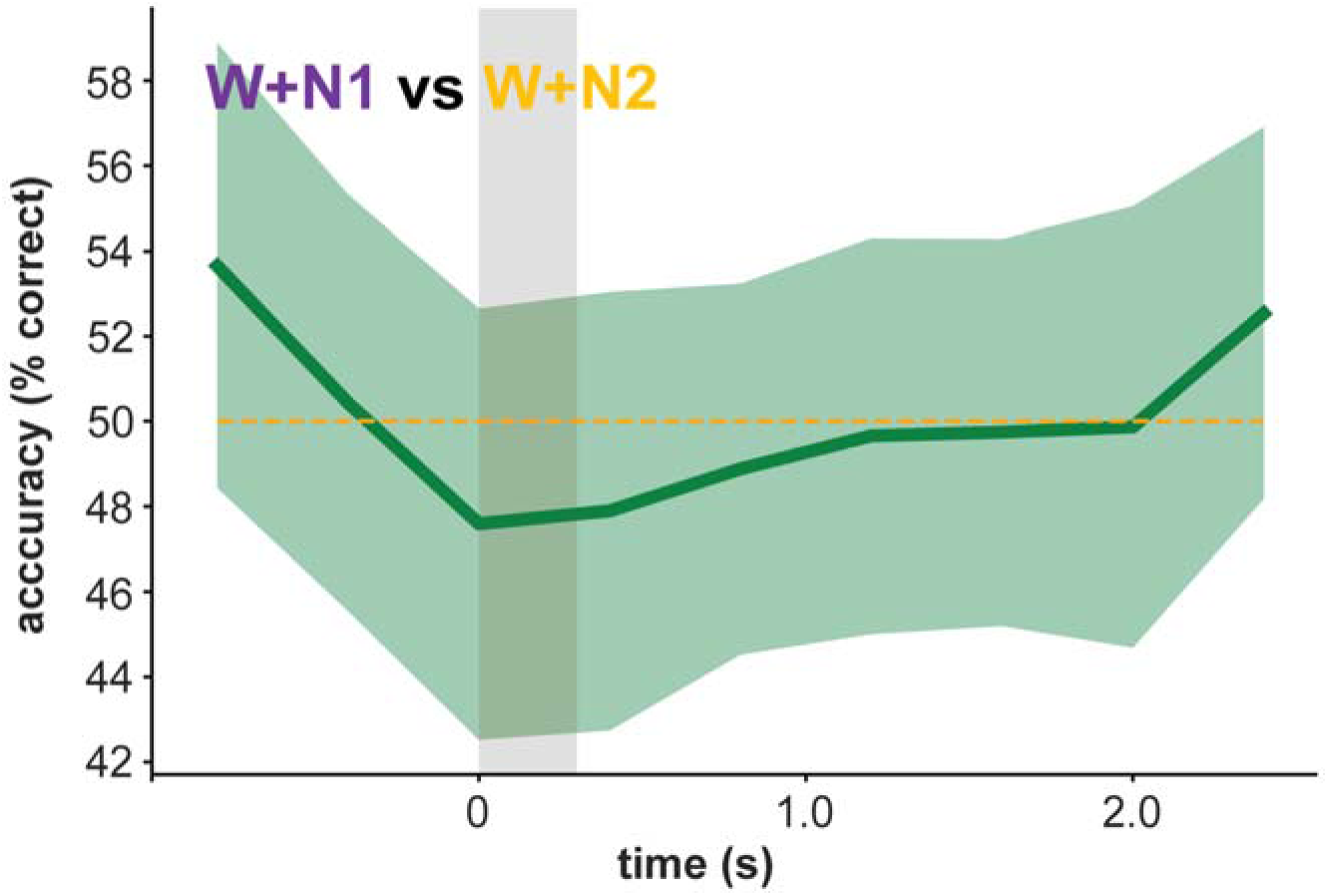
W+N1 vs W+N2 multilayer perceptron performs around chance after reward pairing. Cross-validated W+N1 vs W+N2 MLP accuracy vs. time after pairing both W+N1 and W+N2 with reward. Bin size equals 400 ms. Bold line: mean cross-validated MLP accuracy across 100 pseudosimultaneous sessions generated from 8 mice and 2,926 neurons plus associated neuropil regions-of-interest for each (see methods). Shaded error region: standard deviation across 100 pseudosessions. Shaded rectangle: auditory stimulus epoch. Yellow dashed line: chance performance level.

